# Carbonyl stress primes the metastable aging brain for Alzheimer’s disease

**DOI:** 10.64898/2026.07.24.740487

**Authors:** Aurélien M. Badina, Eléonore Raveloson, Isabel Meister, Quentin Amossé, Laurene Abjean, Kelly Ceyzériat, Stergios Tsartsalis, Serge Rudaz, Philippe Millet, Benjamin B. Tournier

## Abstract

Alzheimer’s disease (AD) is defined by amyloid-β (Aβ) plaques and tau tangles, yet the inflammation that comes with it is only partly localized where those lesions accumulate. Using spatial transcriptomics on 16 human hippocampal sections containing adjacent cortex, we found that plaques concentrated in grey matter, especially cortex. In contrast, the strongest inflammatory response occupied white matter and increased with distance from Aβ-positive spots. This inflammatory signature increased with Braak stage in an independent 31-subject hippocampal bulk proteomic cohort. The white-matter environment revealed a distinct chemistry, with lipidomics showing cortical white matter gaining cholesteryl esters, lysosomal storage lipids and peroxidation-prone polyunsaturated species while losing myelin lipids, alongside carbonyl, glycation and iron-handling signatures. In an external single-nucleus cohort, an oligodendrocyte lipid-droplet program tracked cognitive decline after adjustment for amyloid and tangles, and the same reactive-glia chemistry recurred above expression-matched nulls across seven neurodegenerative datasets. We propose that this lipid-rich glial environment is a metastable, primed state, and that Aβ and tau act as catalysts that tip it toward a self-sustaining inflammatory reaction.

**Highlights:**

- Plaques concentrate in grey matter, whereas inflammatory programs increase with distance and peak in glial/myelin-rich white matter.
- The white-matter response strengthens with Braak stage and is characterized by carbonyl stress, lipid storage, and myelin loss.
- An oligodendrocyte lipid-droplet program tracks cognitive decline after adjustment for Aβ and tau.
- A lipid-droplet reactive-glial state recurs across neurodegenerative diseases with distinct triggers and vulnerable cells.
- Aβ and pTau may tip a metastable tissue environment already primed by lipid and carbonyl stress.

**Graphical abstract:** 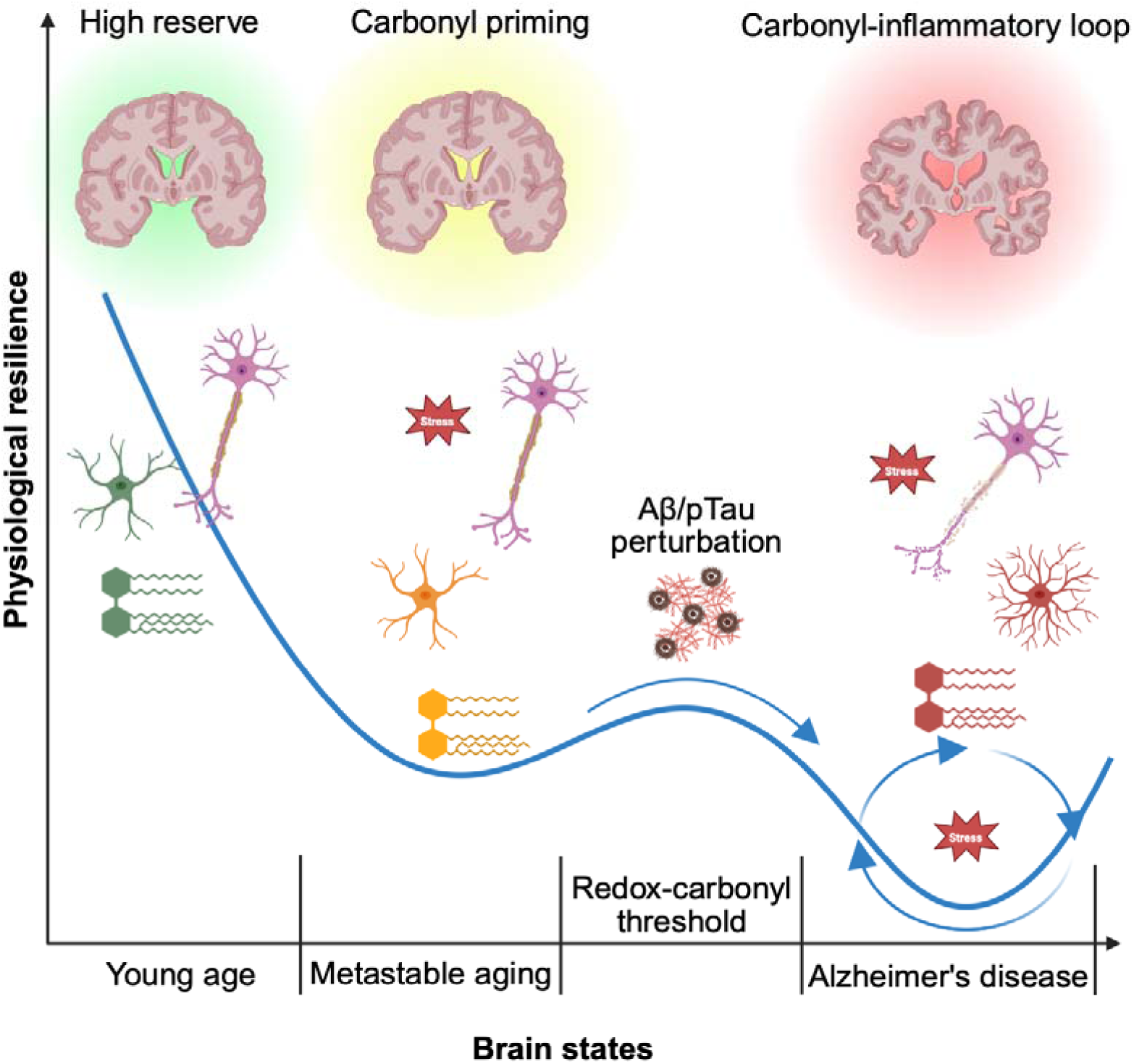

## Introduction

A false vacuum is a metastable system, a system that appears stable while occupying a higher-energy state than its true ground state where a sufficient perturbation can move it from this metastable state toward a lower-energy state. Here, we use this idea as a framework for the aging and pathological brain.

Alzheimer’s disease (AD) is neuropathologically defined by amyloid-beta (Aβ) plaques and hyperphosphorylated tau (pTau) tangles, with severity determined through amyloid, tau and neuritic-plaque lesion accumulation [1–4]. Current models place Aβ, pTau and their interaction as causing much of the inflammatory response [5–14]. Microglia and astrocytes activate around deposits, chemokine networks expand from these lesions, and neuronal injury follows Aβ-or pTau-mediated toxicity. AD inflammation also extends beyond the immediate plaque-proximity response, with vascular dysfunction, white-matter and myelin abnormalities, oxidative stress and plaque-associated glial niches documented brain-wide. Non-demented and resilient subjects can accumulate lesions while retaining cognition, showing that plaque burden alone does not determine the clinical transition to dementia. These observations raise the question of whether part of AD’s inflammatory pathology is simply a response to plaques or an independent mechanism present in the tissue environment, on which plaques and tangles accumulate. In a physiological state, antioxidant defense, glymphatic clearance, membrane-lipid homeostasis and vascular integrity contribute to brain-tissue homeostasis [15, 16]. Over a lifetime, myelin- and lipid-rich, glia-populated tissue accumulates carbonyl species, advanced glycation end products (AGEs), oxidized lipids, iron-handling pressure and clearance stress. On this basis, we propose that Aβ and pTau may act as catalysts on an environment that may already be chemically primed for inflammation.

The brain environment is mostly made up of lipids, between 50 to 55% by dry weight, varying between regions [17]. These lipids undergo continuous changes, the most notable being carbonyl stress. This refers to reactive carbonyl species generated by lipid peroxidation and glycoxidation, modifying membrane lipids and proteins, seeding AGEs formation and activating receptor for advanced glycation end products (RAGE) and nuclear factor-κB (NF-κB) inflammatory signaling in glial and vascular-associated cells [18–22]. They also interact with iron handling and ferroptosis pathways: poly-unsaturated fatty acids (PUFA)-rich membranes are peroxidation substrates, iron catalyzes radical propagation, and each oxidative event generates reactive products that feed the next round [22–33].

These observations led us to ask whether the spatial organization of AD inflammation is determined primarily by local plaque burden or by the biochemical and cellular state of the tissue in which plaques accumulate. We mapped inflammation in human hippocampal formation and adjacent cortex along the plaque-distance axis and across AD stage. Because the hippocampus is allocortical, neocortical grey matter (GM) and white matter (WM) terminology does not fully apply. We therefore use grey-matter-like (GML) for neuronal and soma-rich hippocampal territories, and white-matter-like (WML) for adjacent axonal, myelin-rich and glial territories. We combined spatial transcriptomics with bulk proteomics, lipidomics, cell deconvolution, spatial ligand-receptor modeling, external single-nucleus validation, cross-disease public datasets and immunohistological confirmation to define the inflammation’s anatomy, chemistry, cell-type attribution and temporal trajectory.

## Results

### 1. Inflammation localizes away from plaques, in glial and myelin-rich tissue

A cohort of 16 FFPE human hippocampal sections containing adjacent cortex was scanned and sequenced with Visium technology (CT 7, AD 9; Braak I-VI), with 55 µm spots spaced 100 µm center-to-center and approximately 5,000-7,000 genes detected per spot (Supplemental Data Table S1; section-level spatial quality control and spot composition in Supplemental Data Table S2). We manually annotated Aβ-positive spots and anatomical compartments and investigated gene expression accordingly.

The spatial Aβ burden was higher in AD: CT samples carried 0.88% plaque-positive spots on average, compared to 9.06% in AD samples (**Fig. 1A**; Mann-Whitney two-sided p = 0.0021). We annotated hippocampal GML and WML, surrounding cortical GM and WM compartments and studied plaque accumulation within these compartments using paired per-slide comparisons. In the surrounding cortex, plaques accumulated predominantly in GM in AD (**Fig. 1B**; GM 16.50% versus WM 0.67%; paired p = 0.0039), with the same direction in CT below significance (GM 1.50% versus WM 0.04%; paired p = 0.063). The hippocampus showed the same direction: CT showed no difference in plaque accumulation between GML and WML (**Fig. 1C**; GML 0.82% vs. WML 0.88%, paired p = 1.0), but plaques were higher in GML than WML in AD (6.70% versus 3.16%; paired p = 0.039). Thus, plaque distribution does not differ between compartments in CT, whereas in AD plaques concentrate in the soma- and neuron-rich GML/GM compartment (Supplemental Data Table S3).

To annotate sequencing spots, Aβ plaques were marked by anti-Aβ42 staining on each slide (**Fig. 1D**), and spots were manually classified as Aβ-positive. Genes were analyzed based on their expression on each spot. For visualization, the presynaptic marker synaptosomal-associated protein 25 (SNAP25) marked neuronal/synaptic territories in plaque-rich regions (**Fig. 1E**), myelin basic protein (MBP) marked myelin-rich WML territories (**Fig. 1F**), and glial fibrillary acidic protein (GFAP), a marker of reactive astrogliosis, followed the WML-enriched glial compartment (**Fig. 1G**).

We first investigated the four annotated compartments, cortical GM/WM and hippocampal GML/WML, as biologically distinct tissues. GM and GML were enriched for neurons and synaptic genes and pathways, while WM and WML showed oligodendroglial, astrocytic, microglial, endothelial and extra-cellular matrix (ECM)/inflammatory enrichment (**Fig. 1H**; Supplemental Data Table S4). These data separate compartment identity from plaque burden and indicate that CT WM/WML already carries the glial, myelin-rich and inflammatory/ECM state, while AD plaque deposition takes place in neuronal environment.

**Figure 1.**
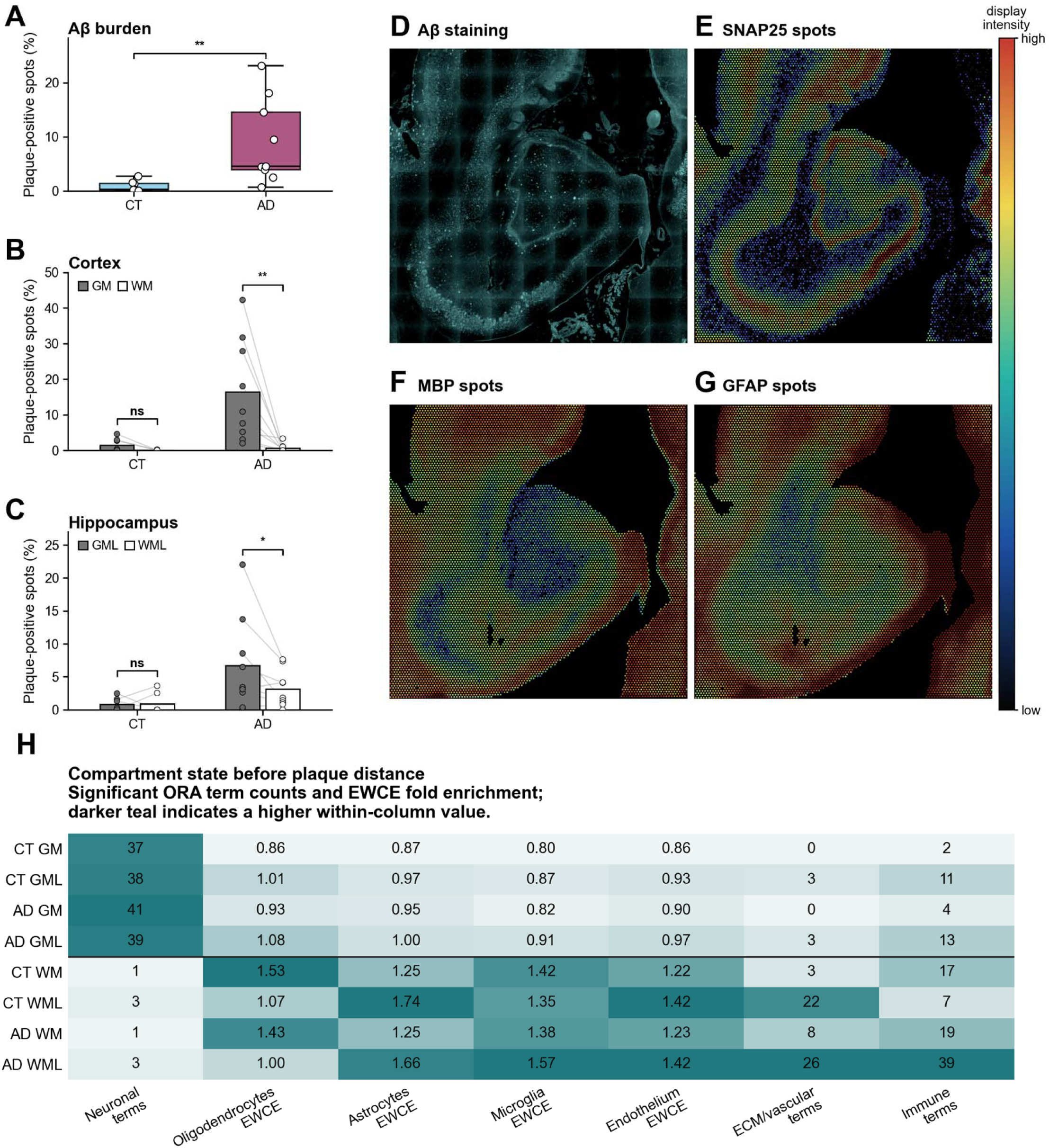
Aβ burden, anatomical marker scaffold and compartment state. **A.** Per-section Aβ burden in CT and AD (percentage of plaque-positive spots from manual spot annotations). Aβ burden was higher in AD than CT sections (CT: 0.88%; AD: 9.06%; two-sided Mann-Whitney p = 0.0021; CT n = 7, AD n = 9). **B.** Paired per-slide plaque density in cortical GM and WM in CT and AD. Cortical plaques were GM-enriched in AD (CT GM: 1.50%, CT WM: 0.04%, paired Wilcoxon p = 0.0625; AD GM: 16.50%, AD WM: 0.67%, paired Wilcoxon p = 0.0039). **C.** Paired per-slide plaque density in hippocampal GML and WML in CT and AD. Hippocampal plaques showed the same direction as cortical densities (CT GML: 0.82%, CT WML: 0.88%, paired Wilcoxon p = 1.000; AD GML: 6.70%, AD WML: 3.16%, paired Wilcoxon p = 0.0391). **D.** Aβ staining on a representative hippocampal/cortical Visium section. **E, F, G.** SNAP25, MBP, and GFAP spot-level signal on the same section, shown with the shared display-intensity scale. **H.** Compartment-state heatmap summarizing compartment-up genes before plaque-distance modeling, with GM/GML rows followed by WM/WML rows. ORA columns show counts of significant GO Biological Process, Reactome, MSigDB Hallmark, and KEGG terms (BH-adjusted p < 0.05) whose descriptions matched neuronal/synaptic, ECM/vascular, or immune/inflammatory themes. EWCE columns show fold enrichment for the indicated reference cell classes; color is scaled within each column.

We then investigated gene expression changes as a function of plaque distance in the hippocampus-centered dataset. In each compartment, spots were pooled into 100µm distance shells per slide before pseudobulk modeling. In the pooled CT+AD model, positive distance coefficients marked genes with higher expression farther from plaques, and negative coefficients marked near-plaque genes (**Fig. 2A**). Per-gene distance coefficients for the CT-only and AD-only models are provided in supplemental (Supplemental Data Tables S5 and S6). This gene-level split separated a glial/stress branch, whose expression increased far from plaques, from a neuronal/synaptic branch, whose expression increased near plaques To investigate possible inflammatory responses in further detail, we manually created a 49 inflammatory gene modules taxonomy covering innate sensing, cytokine signaling, complement, microglial and astrocytic reactive states, AGE-RAGE/NF-κB signaling, ferroptosis/iron handling, BBB/vascular adhesion, glymphatic biology and ECM remodeling (Supplemental Data Table S7; detection coverage in Supplemental Data Table S8). Out of the 49 modules representing various biological functions, 13 modules were significantly overexpressed with plaque distance, which we then pooled into a 116-gene inflammatory gradient core (**Fig. 2B**; Supplemental Data Table S9) to later study the inflammatory functions taking place far from plaques. To visualize how these individual modules behaved across the plaque-distance axis, we plotted representative anchor modules in AD sections using exact 100-µm shells over 0 < distance <= 900 µm, excluding plaque-covered spots and all spots beyond 900 µm (**Fig. 2C**). Of note, the 900µm threshold was chosen to keep the curve within well-sampled shells, with the WML>GML geometry remaining unchanged. The curves showed the same compartmental geometry as the gene-level model with WML module scores being consistently higher than GML scores and generally increasing with distance from plaques, whereas GML curves remained lower, flat, or slightly decreasing. This was visible across microglial/TREM2, disease-associated astrocyte, NF-κB, iron-handling, AQP/glymphatic, and ECM-remodeling modules.

**Figure 2.**
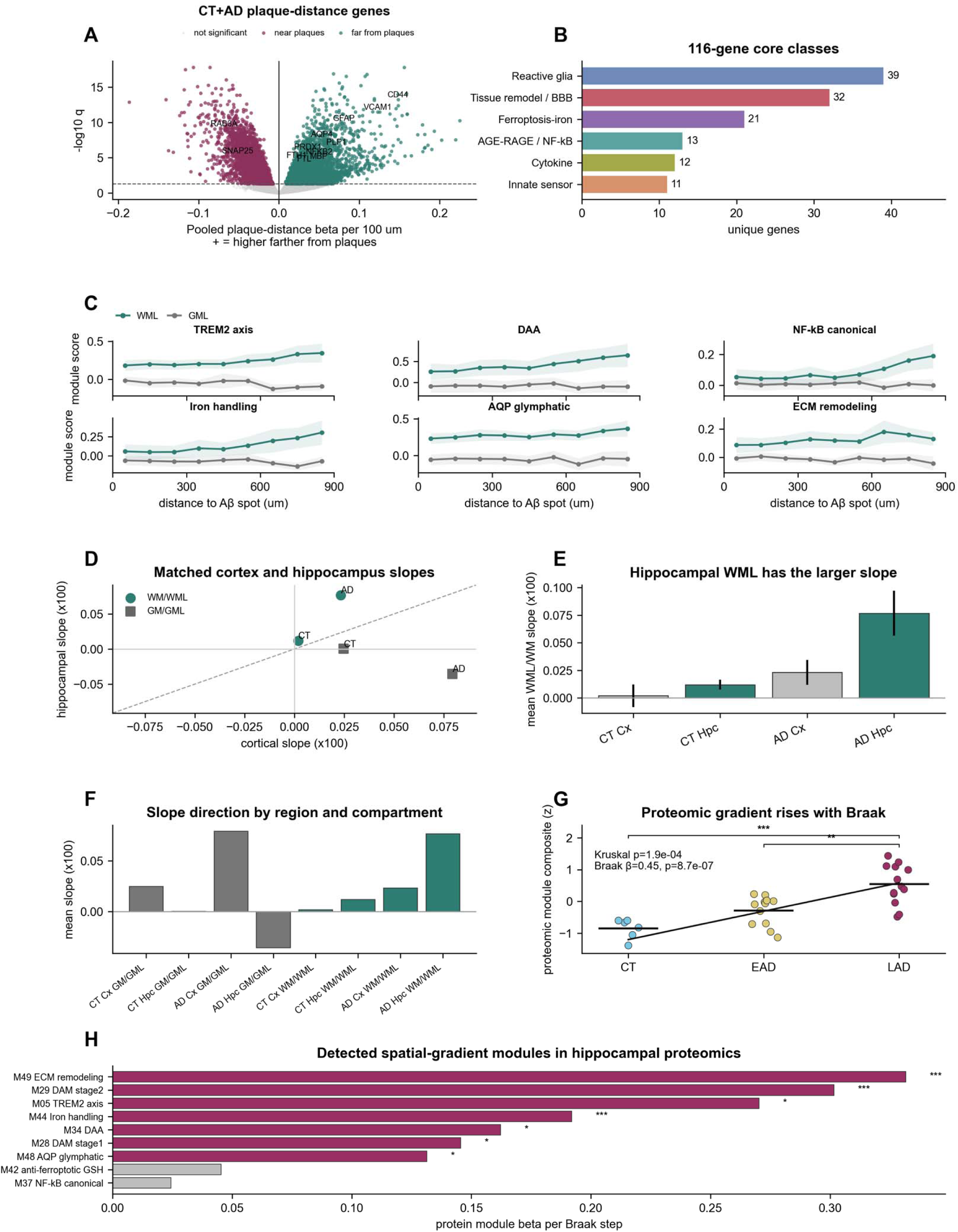
Spatial and temporal properties of the plaque-distance inflammatory gradient. **A.** Gene-level plaque-distance volcano from the pooled CT+AD model. Positive beta indicates higher expression farther from plaques. Of 16,509 tested genes, 3,290 were higher farther from plaques, 3,359 were higher nearer plaques, and 9,860 were not significant at q < 0.05. **B.** Pathway and cell-state classes across the 13 WML-far/GML-near modules whose union defines the 116-gene core: reactive glia, 39 genes; tissue remodel/BBB, 32; ferroptosis-iron, 21; AGE-RAGE/NF-κB, 13; cytokine, 12; innate sensor, 11. **C.** AD distance curves for representative anchor modules, recomputed from exact spot-level distances using 100-µm shells over 0 < distance <= 900 µm. Lines show mean module score across slide-level shell means; shading shows SEM. Aβ-positive spots at distance 0 and all spots >900 µm were excluded. The final 800-900 µm shell retained 555 WML spots across 8 AD slides and 598 GML spots across 7 AD slides. **D.** Matched cortex/hippocampus WM/WML and GM/GML slopes for the inflammatory-gradient program. **E.** Mean WM/WML slopes by diagnosis and region, showing the largest slope in AD hippocampal WML (mean slope = 7.68 × 10^-4 per um; plotted as x100 = 0.0768). **F.** Mean plaque-distance inflammation slopes by region, compartment, and diagnosis. **G.** Proteomic inflammatory-gradient composite across CT, EAD, and LAD hippocampal proteomes (CT n = 6, EAD n = 12, LAD n = 13). Group means were CT -0.85, EAD -0.29, and LAD 0.54 z units; Kruskal-Wallis H = 17.14, p = 1.90 × 10^-4. Pairwise two-sided Mann-Whitney tests with BH correction gave CT versus EAD q = 0.0529, EAD versus LAD q = 0.00318, and CT versus LAD q = 2.21 × 10^-4. The same composite increased with Braak stage by linear regression (beta = 0.445 per Braak stage, p = 8.67 × 10^-7, R² = 0.572). **H.** Module-level Braak progression for the proteomics-detected spatial-gradient modules. Seven of nine displayed modules passed BH q < 0.05; the strongest effects were M49 ECM remodeling (beta = 0.332, q = 2.7 × 10^-5), M29 disease-associated microglia (DAM) stage 2 (beta = 0.302, q = 4.6 × 10^-4), and M44 iron handling (beta = 0.192, q = 4.6 × 10^-4).

The surrounding cortex, where plaques were more abundant, provided the plaque-rich comparator for the hippocampal gradient. In the WM and WML compartments, cortex and hippocampus moved in the same direction: module-score slopes were positive in both regions, and hippocampal WML was larger than cortical WM (**Fig. 2D, E**; paired Wilcoxon p = 0.003). The larger hippocampal WML slope fits a myelin-rich and glia-rich compartment with high circuit, metabolic and clearance demand.

In the GM and GML compartments, the gradient’s geometry differed between cortex and hippocampus. Cortical GM had a positive plaque-distance inflammation slope, whereas hippocampal GML was neutral to negative in AD (**Fig. 2F**; rho = -0.07, p = 0.803). The plaque map explains this opposition, as plaques were mainly GM/GML-biased, whereas the strongest inflammatory tissue response was present in glial/myelin-rich WM/WML.

### 2. The spatial inflammatory gradient shows Braak-progressive carbonyl/redox proteomic programs

To investigate in further details the inflammatory gradient along the evolution of the disease and test whether the spatially detected inflammatory gradient progressed with AD stage, we used an independent 31 subject hippocampal cohort (CT 6, EAD 12, LAD 13; Braak II-VI), on which we performed a bulk proteomics analysis.

Of the 13 inflammatory modules expressed far from plaques, nine had sufficient proteomic coverage. Their per-subject field composite increased with Braak stage (**Fig. 2G**; beta = +0.45 per Braak stage, p = 8.7 × 10^-7), and most individual modules moved in the same direction (**Fig. 2H**).

The same 49-module taxonomy was scored in the proteomics dataset. Twenty modules, including the nine previously detected, were detected in the proteomics dataset, all had positive Braak slopes, and 11 passed BH q < 0.05 (**Fig. 3A**, Supplemental Data Table S10; sensitivity analyses in Supplemental Data Table S11). The composite field score increased across Braak stage in the same subjects (**Fig. 3B**). The strongest WML-gradient anchors were M49 ECM-remodeling, M29 DAM stage 2, M44 iron-handling, M34 disease-associated astrocyte and M48 AQP/glymphatic modules (**Fig. 3C**).

Because each Visium spot mixes several cell types, a module score alone does not say which cells carry the signal. To assign the gradient modules to cell types, we correlated each per-(slide × distance-shell × compartment) module score with Robust Cell-Type Decomposition (RCTD) cell-type weights across the full broad-class matrix (Supplemental Data Table S12; recovery of expected module cell origins in Supplemental Data Table S13). The main classes are shown in **Fig. 3D, E**. The anchor modules correlated strongest with astrocytic, microglial or vascular-associated weights, defining a mixed-spot attribution layer.

**Figure 3.**
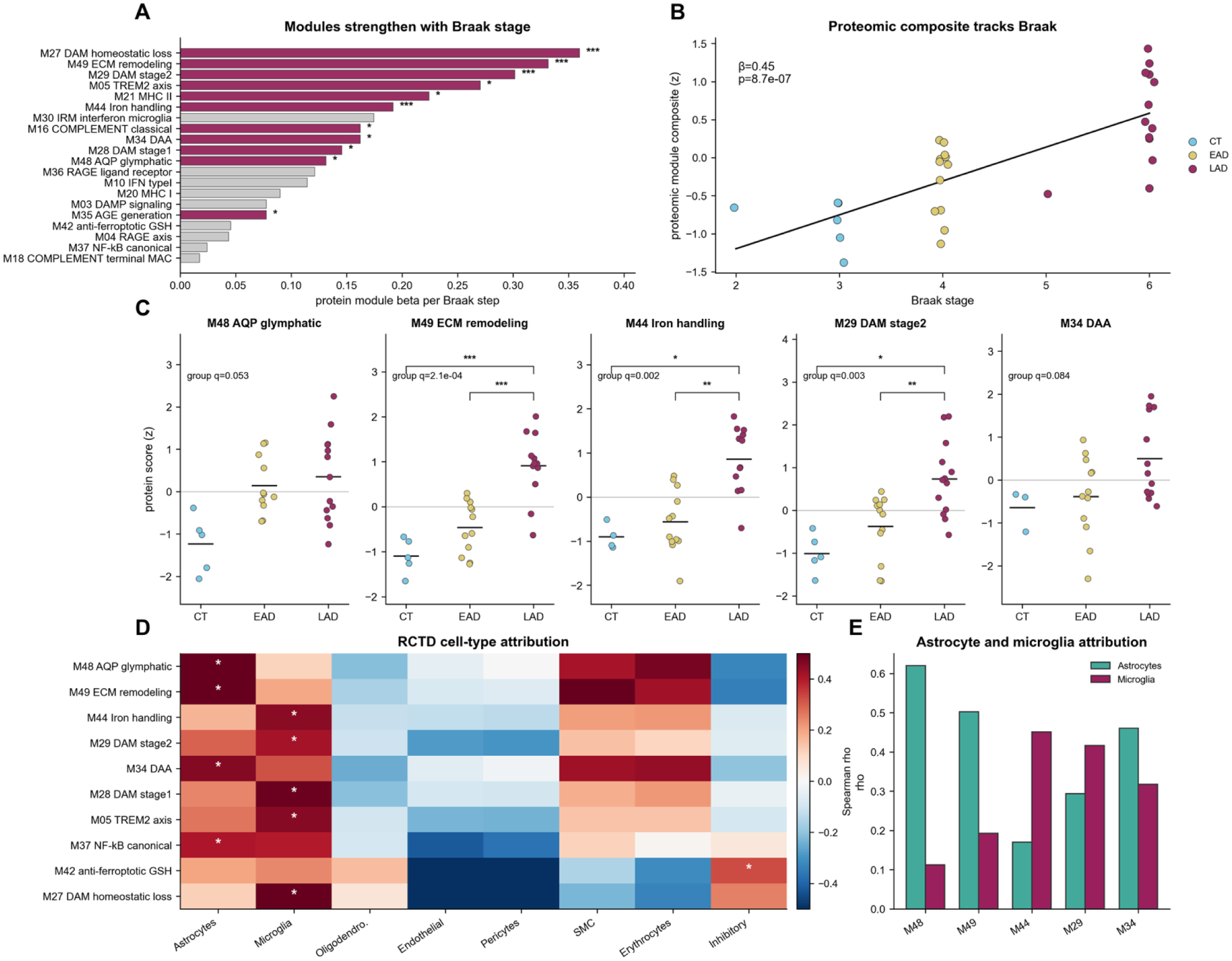
Module-level temporal progression and cell-type attribution. **A.** Braak slopes across the 20 proteomics-testable inflammation modules in hippocampal bulk proteomics. All 20 modules had positive Braak slopes and 11 passed BH q < 0.05; stars indicate BH-corrected Braak significance. The largest effects were observed for DAM homeostatic loss (β = 0.360, q = 5.0 × 10^-4), ECM remodeling (β = 0.332, q = 2.7 × 10^-5), DAM stage 2 (β = 0.302, q = 4.6 × 10^-4), TREM2 axis (β = 0.270, q = 0.026) and MHC-II (β = 0.224, q = 0.015). **B.** Proteomic inflammatory-gradient composite across Braak stages in 31 subjects. Linear regression gave β = 0.445 z units per Braak stage, p = 8.67 × 10^-7, R² = 0.572; Spearman ρ = 0.780, p = 2.35 × 10^-7. **C.** Stage trajectories for five anchor modules across CT, EAD and LAD. Points are subjects and bars are group means; group q-values and BH-corrected pairwise comparisons are shown. ECM remodeling (group q = 2.08 × 10^-4), iron handling (q = 0.00165) and DAM stage 2 (q = 0.00322) showed significant group progression; AQP/glymphatic was borderline after correction (q = 0.0529) and DAA did not pass group-level correction (q = 0.0837). **D.** Selected broad-class RCTD cell-type attribution map using the Green et al. 2024 ROSMAP DLPFC reference. Stars mark the strongest positive association per module. **E.** Astrocyte and microglia attribution for the five glial anchor modules, showing astrocyte-biased attribution for M48 AQP/glymphatic (ρ = 0.620), M49 ECM remodeling (ρ = 0.502) and M34 DAA (ρ = 0.461), and microglia-biased attribution for M44 iron handling (ρ = 0.451) and M29 DAM stage 2 (ρ = 0.416). The full correlation matrix is provided in Supplemental Data Table S12.

As AGE-RAGE, carbonyl and oxidative stress pathways and responses were repeatedly present in the data as being overexpressed far from plaques and with a stronger expression through AD stages, we manually established a deeper 31-gene carbonyl-stress core (Supplemental Data Table S14). We defined it from genes with direct roles in methylglyoxal/glyoxalase detoxification, fructosamine repair, aldo-keto/aldehyde carbonyl clearance, NRF2-glutathione antioxidant response and peroxide/lipid-peroxidation detoxification. This carbonyl module directly followed the spatial far-from-plaques gradient and increased in hippocampal proteomics across AD stage (**Fig. 4A, B**). Six carbonyl/redox submodules resolved the chemistry inside this aggregate score: MGO/glyoxalase detoxification, aldehyde-carbonyl clearance, AGE-RAGE ligand load, NRF2/glutathione redox defense, lipid-peroxidation/ferroptosis and protein-damage/proteostasis (Supplemental Data Table S14). WML exceeded WM in CT and AD across all six signatures, with paired WML-WM differences strongest for glutathione/NRF2, protein-damage/proteostasis and aldehyde-carbonyl clearance (**Fig. 4C**).

Single-protein analyses pointed in the same direction (**Fig. 4D**). In the bulk proteome, 944 proteins reached p < 10^-3 and |r| > 0.5 against the proteomic inflammatory gradient (Supplemental Data Table S15). Selected carbonyl-axis or AGE-RAGE-aligned correlates included AHNAK (r = +0.91), MSN (+0.91), PRDX1 (+0.89), CD44 (+0.89), GFAP (+0.81), NQO1 (+0.75), PRDX6 (+0.73) and MAOB (+0.71). Because single-protein correlations test proteomic evidence alone, we complemented them with a broader gene-level audit that required convergent spatial plaque-distance and proteomic-Braak evidence for the same gene, providing a cross-modal specificity. This audit recovered the same polarity and is reported as a supplemental robustness screen (Supplemental Data Table S16).

Because WML also carried endothelial and perivascular signatures, we tested whether this inflammatory gene set could simply be due to blood-brain barrier (BBB) dysfunction causing leakage or peripheral immune infiltration. The vascular signatures were themselves higher in hippocampal WML than GML. Even so, the WML inflammatory-glial plaque-distance effect survived joint adjustment for all three vascular scores (β = +0.038, p = 0.0037), as did the carbonyl/oxidative score (β = +0.032, p = 6.6 × 10^-8). The inflammatory and carbonyl gradients are therefore not explained by BBB dysfunction, vascular signal or blood contamination (Supplemental Data Table S17).

**Figure 4.**
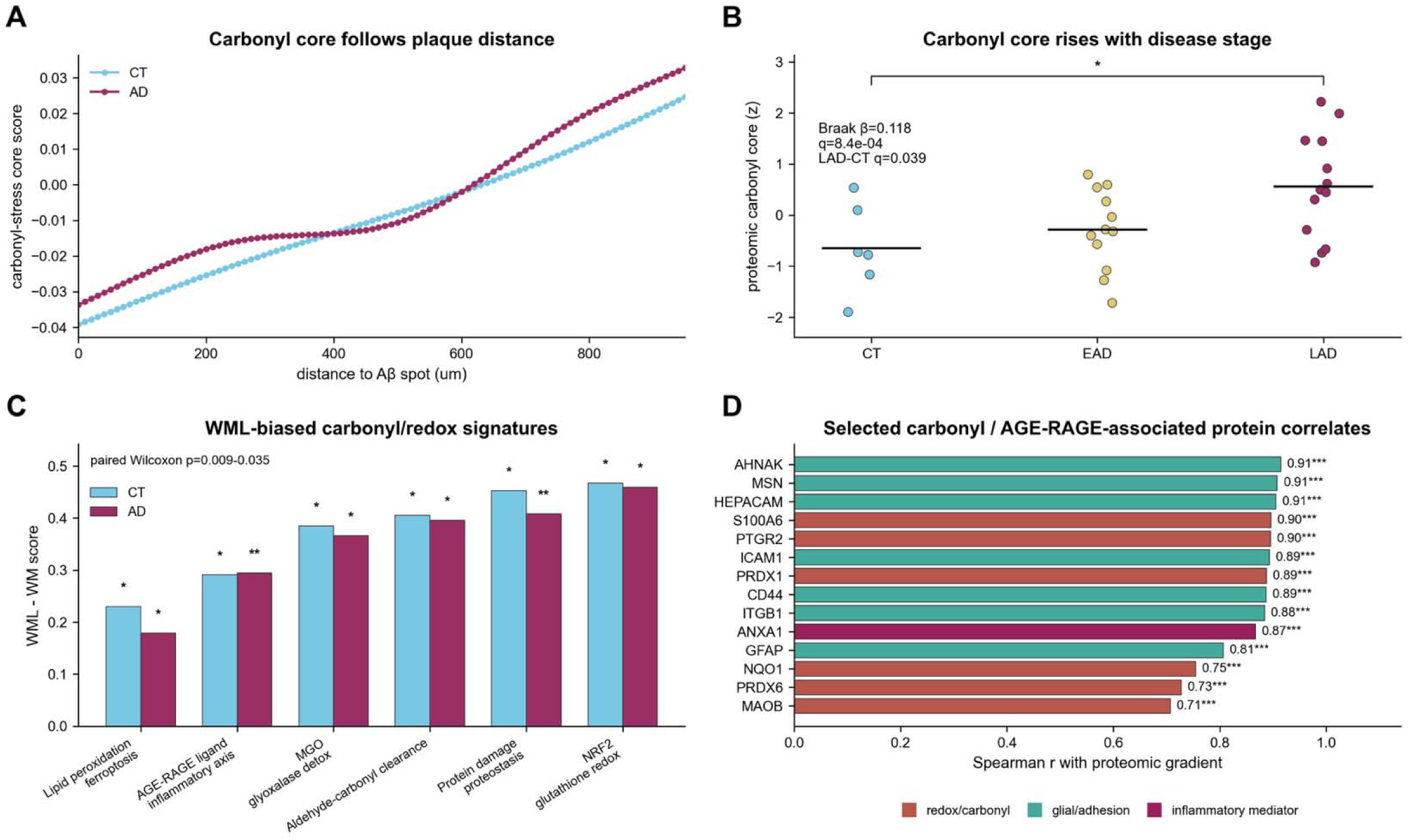
AGE-RAGE, carbonyl and oxidative-stress evidence. **A.** Carbonyl-stress core spatial distance curve in CT and AD, displayed up to 950 µm from Aβ-positive spots. **B.** Carbonyl-stress core per-subject proteomic abundance by stage. The Braak-stage model gave β = 0.118, q = 8.39 × 10^-4. The LAD-versus-CT contrast was significant (q = 0.0392), whereas EAD-versus-CT was not significant (q = 0.309). **C.** Paired WML-WM compartment contrasts for curated carbonyl/redox-handling signatures. Positive values indicate WML-biased signal, paired Wilcoxon: CT n = 7, p = 0.022-0.035; AD n = 9, p = 0.009-0.024. **D.** Selected carbonyl/AGE-RAGE-associated protein correlates ranked by Spearman correlation against the proteomic inflammatory gradient and colored by biological class. Displayed correlations ranged from MAOB (r = 0.707, q = 1.39 × 10^-4 to AHNAK (r = 0.915, q = 1.69 × 10^-9; all displayed proteins passed q < 0.001.

### 3. Lipidomics identifies the chemical environment behind the glial/myelin-rich state

The spatial and proteomic data both pointed to a lipid-rich glial and myelin environment defined at the level of mRNA and protein, implicating carbonyl stress responses. We therefore measured that environment directly by profiling lipids in an independent paired grey and white human-tissue cohort of 90 samples from 49 subjects (cortex CT 6, EAD 11, LAD 13, Braak II–VI; hippocampus CT 11, EAD 2, LAD 6, Braak 0–VI). Cortical samples were separated into grey (GM) and white (WM) matter and hippocampal samples into their grey-matter-like (GML) and white-matter-like (WML) counterparts. As tissue provenance and region were determined by the dissection, we tested every lipid class within region and compartment using compositional CLR scores, with cohort details in STAR Methods.

We first tested white vs. grey matter differences (**Fig. 5A**). In the 11 CT hippocampal pairs, white matter was enriched over adjacent grey matter for myelin glycosphingolipids (median CLR difference +0.78, q = 0.0021), sulfatides (+0.51, q = 0.0021), sphingomyelin (+0.35, q = 0.0021) and ether phospholipids (+0.21, q = 0.0216), and was depleted of cholesteryl esters (−1.14, q = 0.0021), polyunsaturated and highly unsaturated phospholipids (−1.09 and −1.26, q = 0.0021) and triacylglycerols (−0.82, q = 0.0021). The six CT cortical pairs followed the same direction, without reaching corrected significance at this sample size (Supplemental Data Table S18).

The lipidome changed most with disease stage in the cortical WM. Within that compartment, myelin-related lipids decreased while storage and disposal lipids increased (**Fig. 5B**). Cholesteryl esters, the storage form of cholesterol, increased by +1.64 CLR units already in early AD and by +1.70 in late AD relative to control (p = 0.0016, Braak ρ = +0.52, p = 0.0071). Lysosomal bis(monoacylglycero)phosphate (BMP), a marker of the late-endosomal and lysosomal disposal compartment, increased by +0.42 in both early and late AD (p = 0.0048, ρ = +0.49, p = 0.0119). Polyunsaturated fatty acid (PUFA)-containing acylcarnitines, the mitochondrial transport form of fatty acids entering β-oxidation, increased from +0.49 in early AD to +0.63 in late AD (p = 0.0120, ρ = +0.44, p = 0.0238), together with highly unsaturated phospholipids (ρ = +0.39). The median myelin-glycosphingolipid score decreased by 0.37 CLR units in late AD compared to CT across the same tissue (p = 0.036). Most of these changes were already present in the early-AD group. At the lipid species level, 150 individual lipids separated late-AD from control cortical white matter at q < 0.1, while cortical grey and both hippocampal compartments yielded none at this threshold and sample size (Supplemental Data Table S19).

The accumulated lipid species were polyunsaturated, the potential substrate for oxidation and carbonylation. Among the increased species were docosahexaenoate-carrying cholesteryl ester CE 22:6 and CE 18:2, a ten-double-bond BMP 42:10, and the polyunsaturated acylcarnitines CAR 20:2 and CAR 24:4 (all q < 0.1, Braak ρ between +0.44 and +0.58). The chain-length breakdown localized the acylcarnitine increase to the polyunsaturated species (DB ≥ 2, delta = +0.63) and not to total or long-chain saturated acylcarnitines (delta = +0.03 and +0.30, both n.s.). Among the decreased species were all twenty acyl-hexosylceramide species, the bulk of hexosylceramides, sulfatides, lysophosphatidylcholines, and ether phospholipids fell on balance, with 12 of 17 significant ether-phosphatidylethanolamine species reduced. The ether phospholipids that decreased are endogenous radical sinks, with their loss accompanying the accumulation of oxidizable material.

The increase in storage and lysosomal lipids extended into cortical grey matter, while the myelin and PUFA remodeling stayed specific to white matter. Cortical grey matter gained cholesteryl esters (Braak ρ = +0.57, p = 0.0017), neutral storage lipids (ρ = +0.46, p = 0.0145) and BMP (ρ = +0.45, p = 0.0165) with stage, but showed no myelin glycosphingolipid or PUFA-phospholipid trend. Hippocampal WML also showed the lysosomal BMP increase (ρ = +0.52, p = 0.0344), with four late-AD subjects in that subgroup.

The same lipid-handling axis appeared in the spatial and proteomic data measured in the other cohorts. In the spatial transcriptomics, genes for phospholipid PUFA release (PLA2G4A), lipid-droplet storage (PLIN3), cholesteryl ester synthesis (SOAT1), lysosomal lipid catabolism (NPC2, LIPA), iron storage (FTL, FTH1) and carbonyl and peroxide handling (ALDH1A1, PRDX6, MGST1, CAT) were all higher farther from plaques in both control and AD tissue (ALDH1A1 q = 2.9 ×10^-6, PLA2G4A q = 5.4 × 10^-4, NPC2 q = 6.4 × 10^-5), while the ferroptosis-limiting GPX4 was the one gene higher near plaques (**Fig. 5C**; Supplemental Data Table S20). In hippocampal proteomics, the droplet and lysosomal proteins of the same axis increased with Braak stage: the droplet-associated apolipoproteins CLU (ρ = +0.73) and APOE (ρ = +0.52) [34], the droplet coat protein PLIN3 (ρ = +0.50) and the lysosomal hydrolases NPC2 (ρ = +0.42), GLA (ρ = +0.39) and LIPA (ρ = +0.37), while the PUFA acyl-CoA ligase ACSL4 fell (ρ = −0.50) (**Fig. 5C**; Supplemental Data Table S21). Across the 19 subjects measured on both platforms, per-subject cortex-white acylcarnitine load correlated with the reduced proteomic loop score (ρ = +0.59, p = 0.008; **Fig. 5D**; Supplemental Data Table S22). Lipid-droplet storage, lysosomal disposal, PUFA mobilization and iron and redox pressure move in the same direction in three separate cohorts and three measurement types.

This lipid environment supplies the substrate for white-matter carbonyl and redox chemistry. The cortical white-matter compartment accumulates polyunsaturated cholesteryl esters, BMP and PUFA acylcarnitines, loses its myelin glycosphingolipids, depletes its ether radical sinks, and the carbonyl and redox signatures increased in that same white-matter compartment. Polyunsaturated acyl chains are the direct substrate for lipid peroxidation and for the reactive carbonyls that peroxidation generates, and their storage in droplets and lysosomes alongside a decreasing antioxidant-lipid buffer describes an environment primed for stress.

**Figure 5.**
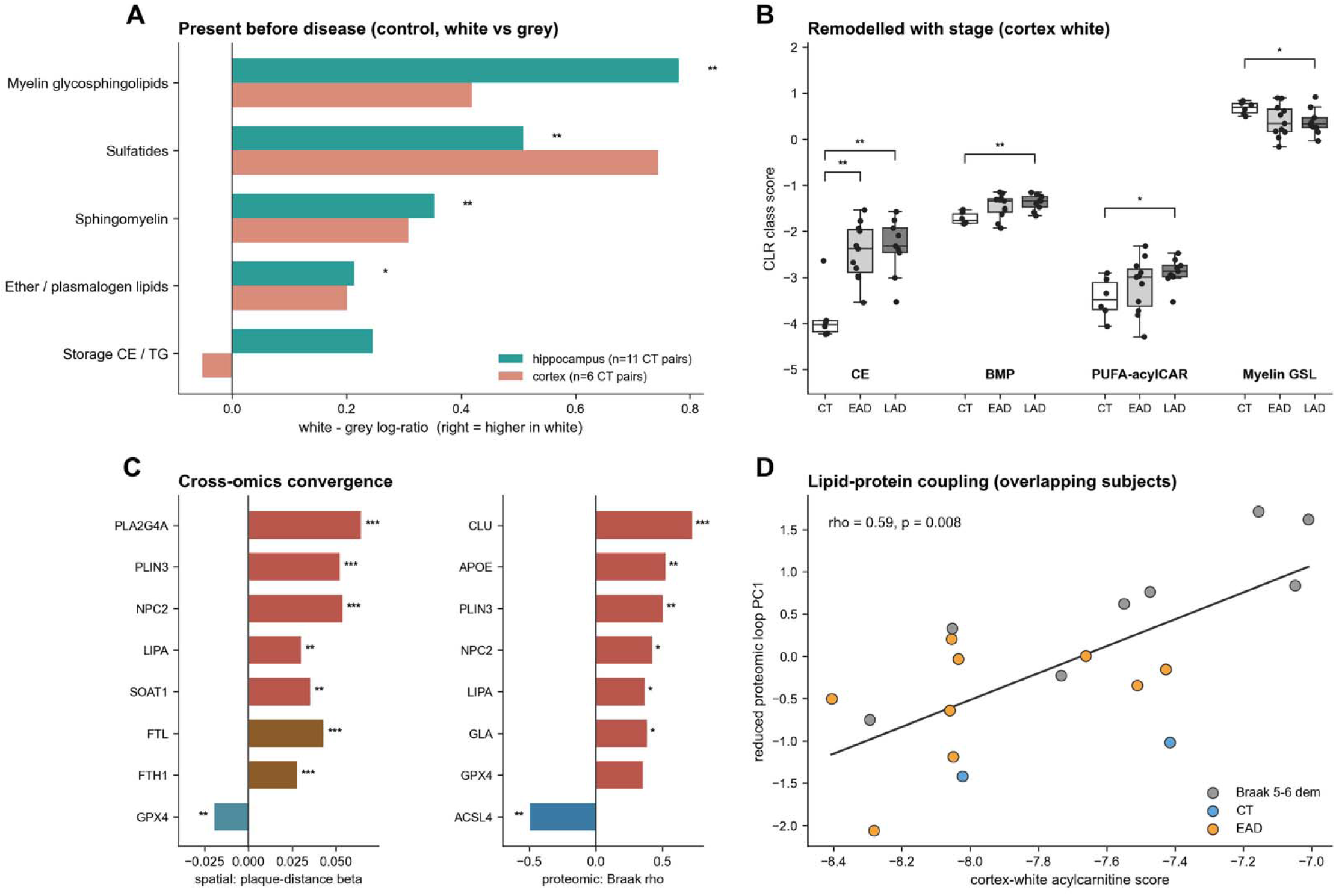
An aged white-matter lipid environment, present before disease and remodeled with stage. **A.** Paired white-minus-grey lipid-class log-ratios in control tissue for hippocampus and cortex; positive values indicate enrichment in white matter. Stars mark BH-significant paired white-versus-grey differences. Hippocampal control donor pairs showed significant enrichment for myelin glycosphingolipids, sulfatides and sphingomyelin (all q = 0.0021), and ether lipids (q = 0.0216; hippocampus n = 11). Cortex control pairs are shown in parallel (n = 6). **B.** Cortex-white CLR class scores across CT, EAD and LAD for cholesteryl esters (CE), lysosomal BMP, PUFA acylcarnitines and myelin glycosphingolipids. Brackets show significant two-sided Mann-Whitney contrasts: CE CT-EAD p = 0.0031 and CT-LAD p = 0.0016; BMP CT-LAD p = 0.0048; PUFA acylcarnitines CT-LAD p = 0.012; myelin glycosphingolipids CT-LAD p = 0.036. **C.** Cross-omics convergence: spatial plaque-distance beta in CT for lipid and redox genes (left) and hippocampal proteomic Braak correlations for lipid-handling proteins (right). Spatial CT effects included PLA2G4A β = 0.064, PLIN3 β = 0.052, NPC2 β = 0.054, FTL β = 0.043 and GPX4 β = -0.020. Proteomic Braak correlations included CLU ρ = 0.725, APOE ρ = 0.524, PLIN3 ρ = 0.504 and ACSL4 ρ = -0.496. **D.** Cortex-white acylcarnitine score versus the reduced proteomic loop PC1 in overlapping subjects (n = 19; Spearman ρ = 0.588, p = 0.0081). Lipid species-level remodeling and the full cell-state evidence matrix are provided in the supplement (Supplementary Fig. 8; Supplemental Data Tables S18-S22).

### 4. The environment response tracks cognitive decline independently of amyloid

After locating and characterizing the inflammatory response, we investigated whether the compartment split between plaque-bearing GML and inflammation-bearing WML was reproduced at the protein level and whether the associated environment and inflammation programs were correlated with clinical symptoms. We therefore used two formalin-fixed paraffin-embedded (FFPE) immunohistochemistry (IHC) resources: a paired spatial-cohort subset stained for AT8 phosphorylated tau, 4G8 Aβ, glial fibrillary acidic protein (GFAP) and ionized calcium-binding adaptor molecule 1 (IBA1) across matched GML/WML zones, and an independent diagnostic series spanning young, CT Aβ−, CT Aβ+/NDAN-like, early-AD and late-AD brains; cohort composition, marker-specific availability and technical exclusions are reported in Methods and Supplemental Data Table S1.

At the protein level, IHC reproduced the plaque-versus-inflammatory environment split that our spatial data showed. We quantified AT8 phospho-tau, 4G8 Aβ, GFAP and (IBA1) by 3,3′-diaminobenzidine (DAB) immunohistochemistry across the harmonized spatial-cohort FFPE sections (**Fig. 6A**). AT8 phospho-tau did not differ significantly between the grey- and white-matter-like zones in either group (**Fig. 6B**; AD 3.78% versus 1.34%, paired p = 0.0645; CT 0.21% versus 0.07%, paired p = 0.0625). Aβ concentrated in the grey-matter-like zone in AD (**Fig. 6C**; 9 of 10 subjects, paired Wilcoxon p = 0.0195), placing the plaque burden in the same GM/GML compartments the spatial lesion map identified. Astrocytic GFAP showed the opposite bias, higher in the white-matter-like zone than the grey-matter-like zone in all 10 AD subjects (**Fig. 6D**; p = 0.0020) and in the same direction in 4 of 5 controls. Microglial IBA1 did not differ significantly between compartments in either group (**Fig. 6E**; AD 4.56% versus 5.12%, paired p = 0.557; CT 18.53% versus 25.61%, paired p = 0.625). The plaque signal and the astrocytic response therefore occupy different compartments in the same sections (**Fig. 6F**), which replicates the spatially detected results (Supplemental Data Table S23).

We next asked whether the environment programs predict how fast people decline, which requires the longitudinal cognitive trajectories our own tissue does not have. To this end, we retrieved a publicly available single-nucleus dataset, the Green et al. 2024 Religious Orders Study and Memory and Aging Project (ROSMAP) dorsolateral prefrontal cortex (DLPFC) atlas, which pairs single-nucleus profiles with donor-level cognitive-decline metadata [13]. We scored the 49 inflammation modules on this dataset, using its published 45,577-nucleus geometric sketch, a geometry-preserving subsample that keeps rare cell states in proportion, and aggregated module scores per donor and cell type across all 424 donors with clinical metadata [13]. For each module and cell type we fit a cognitive-decline model adjusted for age, sex, amyloid and tangles, so that any association is what remains after the two canonical pathologies are accounted for.

**Figure 6.**
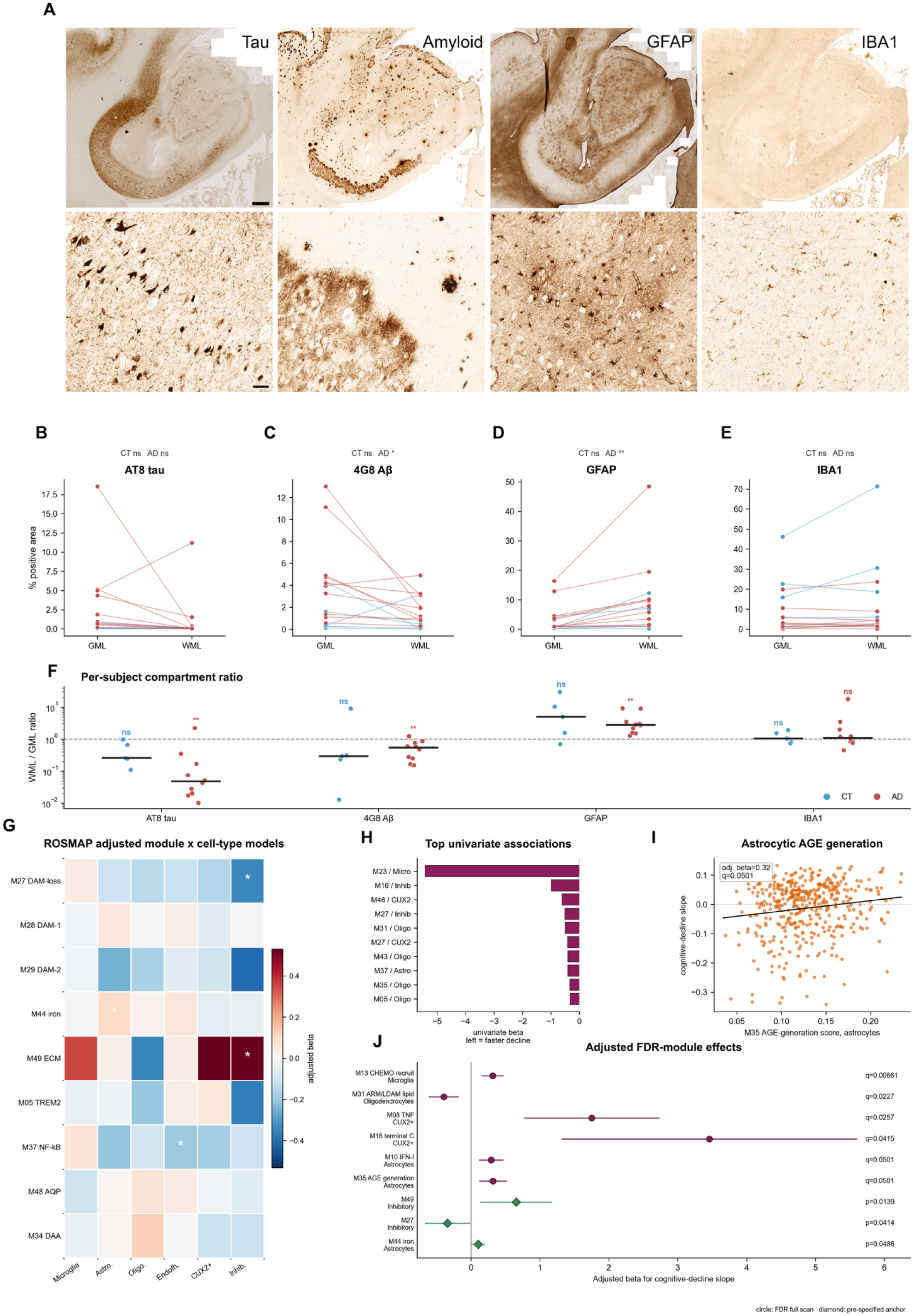
Immunohistological compartment validation and ROSMAP single-nucleus support. **A.** Representative FFPE DAB staining for AT8 tau, 4G8 Aβ, GFAP and IBA1. **B, C, D, E.** Paired GML/WML quantification in the spatial validation cohort (CT n = 5, AD n = 10). AT8 tau was not significantly different between GML and WML in CT (0.21% versus 0.07%, paired p = 0.0625) or AD (3.78% versus 1.34%, paired p = 0.0645). 4G8 Aβ was GML-biased in AD (4.82% versus 1.92%, paired p = 0.0195), but not CT (1.32% versus 0.76%, paired p = 0.4375). GFAP was WML-biased in AD (4.51% versus 11.73%, paired p = 0.00195), but not CT (1.08% versus 4.40%, paired p = 0.125). IBA1 did not differ significantly between compartments in CT (18.53% versus 25.61%, paired p = 0.625) or AD (4.56% versus 5.12%, paired p = 0.557). **F.** Per-subject WML/GML ratios. Median CT/AD ratios were 0.256/0.0475 for AT8 tau, 0.293/0.537 for 4G8 Aβ, 4.94/2.81 for GFAP and 1.04/1.08 for IBA1; AD ratios differed from 1 for AT8 tau (p = 0.00391), 4G8 Aβ (p = 0.00586) and GFAP (p = 0.00195), but not IBA1 (p = 0.492). **G.** ROSMAP adjusted module-by-cell-type models using the Green et al. DLPFC reference; stars mark nominal adjusted-model p < 0.05 in the prespecified anchor display. **H.** Top univariate module-by-cell-type associations with cognitive-decline slope; the strongest was M23/microglia (univariate beta = -5.43, p = 0.00857; amyloid/tangle-adjusted beta = -5.17, p = 0.00403, q = 0.0987). **I.** Astrocytic M35 AGE-generation score versus cognitive-decline slope in ROSMAP (n = 424; Spearman rho = 0.123, p = 0.0111; amyloid/tangle-adjusted beta = 0.318, p = 0.00205, q = 0.0501). **J.** Adjusted FDR-module effects: M13/microglia beta = 0.315, q = 0.00661; M31/oligodendrocytes beta = -0.396, q = 0.0227; M08/CUX2+ beta = 1.75, q = 0.0257; M18/CUX2+ beta = 3.46, q = 0.0415; M10/astrocytes beta = 0.292, q = 0.0501; M35/astrocytes beta = 0.318, q = 0.0501. Prespecified anchor effects are shown for M49/inhibitory neurons (beta = 0.655, p = 0.0139), M27/inhibitory neurons (beta = -0.344, p = 0.0414) and M44/astrocytes (beta = 0.103, p = 0.0486).

The oligodendrocyte lipid-droplet program gave the clearest result and links to the lipid-droplet accumulation measured in the white-matter lipidome. Higher activity of the lipid-droplet-accumulating microglia (LDAM) program (module M31), scored here in oligodendrocytes, tracked faster cognitive decline after amyloid and tangle adjustment (beta = −0.40, q = 0.02), and the association was specific to oligodendrocytes, with microglia, astrocytes and neurons all non-significant for the same program (**Fig. 6G, J**). This is the same lipid-droplet axis that increased with Braak stage in the cortex-white lipidome and in the hippocampal proteome, now carrying information about the rate of decline in an independent cohort, in the white-matter cell type that handles those lipids [35–37]. Three further programs reached the false discovery rate (FDR) threshold in specific cell types: microglial chemokine recruitment (M13, beta = +0.31, q = 0.007), and, in CUX2+ excitatory neurons, tumor necrosis factor (TNF) pathway (M08, q = 0.03) and terminal-complement activity (M18, q = 0.04). Astrocytic AGE-generation activity followed the same pattern (**Fig. 6I**; M35, q = 0.05).

The single-nucleus analysis links the environment to decline but does not order the environment relative to amyloid. To separate the two, we used an independent 54-subject FFPE diagnostic series that includes control brains carrying amyloid without dementia, the NDAN-like state, alongside young, amyloid-negative control, early-AD and late-AD groups [38–40]. Microglial IBA1 and astrocytic GFAP differed across the diagnostic groups (Kruskal-Wallis p = 0.0013 and 0.015; **Fig. 7A, B**), as did the lipid-peroxidation-derived carbonyl 4-hydroxynonenal (4-HNE) (p = 0.0064; **Fig. 7C**). The informative contrast is the amyloid-bearing non-demented brains against AD. Microglial IBA1 in these brains was as low as in amyloid-negative controls and well below early AD (Cliff’s delta = −0.89 versus early AD, q = 0.007), so the microglial inflammatory state was absent in tissue that already carried plaques. Measuring 4-HNE in this same series links the readout to the white-matter carbonyl chemistry and its peroxidation-susceptible lipid substrate, in a cohort where amyloid is present but the response is not yet engaged (Supplemental Data Table S24) [18, 19, 22].

**Figure 7.**
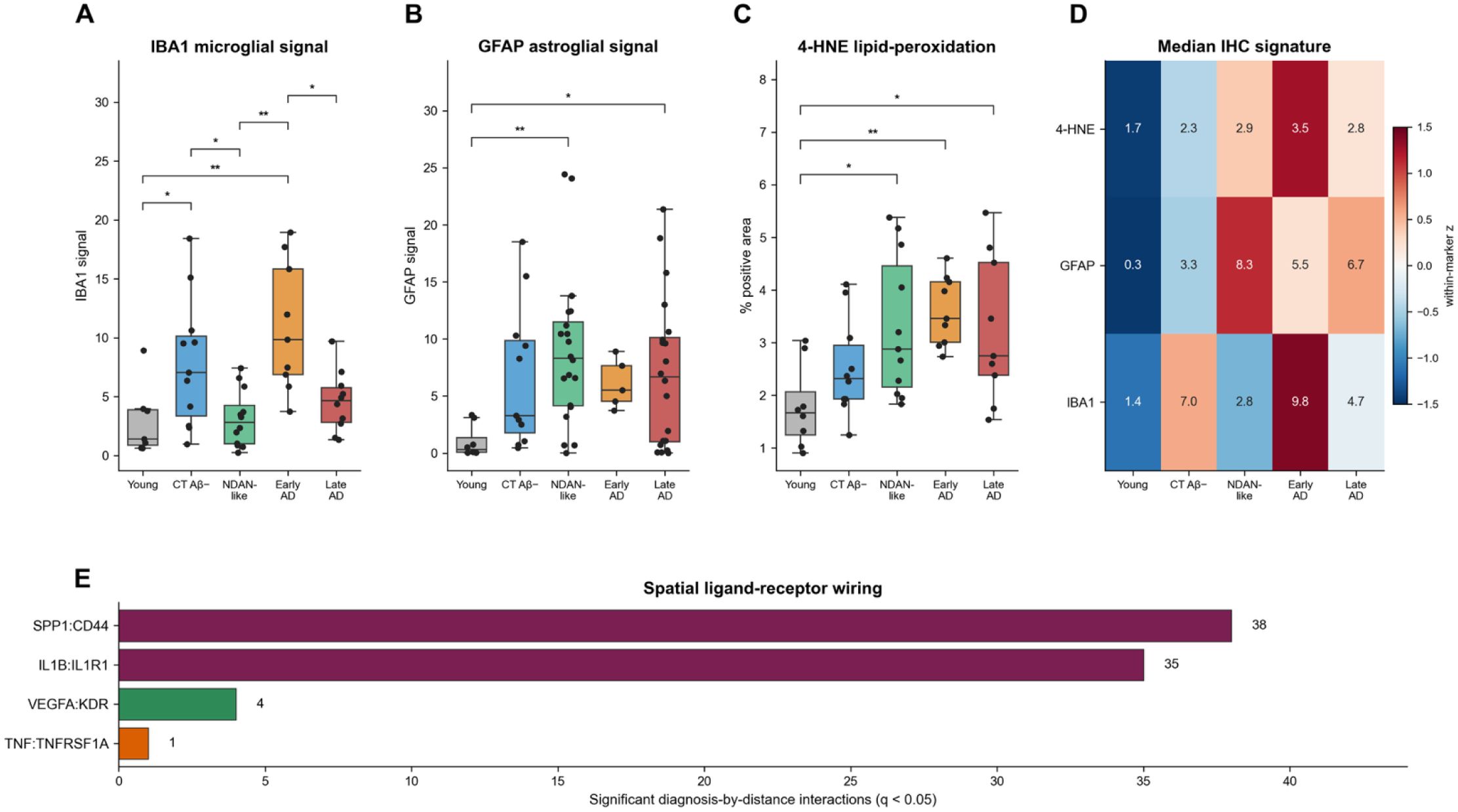
NDAN-like immunohistological series and spatial ligand-receptor wiring. **A, B, C.** IBA1, GFAP and 4-HNE immunohistological signals across Young, CT Aβ−, CT Aβ+/NDAN-like, Early AD and Late AD groups. Kruskal-Wallis tests were significant for IBA1 (n = 49, H = 17.97, p = 0.00125), GFAP (n = 64, H = 12.30, p = 0.0152) and 4-HNE (n = 47, H = 14.30, p = 0.00641). Stars mark Dunn post-hoc tests with BH correction. Significant IBA1 contrasts were Young versus CT Aβ− (pBH = 0.0465), Young versus Early AD (pBH = 0.00544), CT Aβ− versus CT Aβ+/NDAN-like (pBH = 0.0465), CT Aβ+/NDAN-like versus Early AD (pBH = 0.00374) and Early AD versus Late AD (pBH = 0.0465). Significant GFAP contrasts were Young versus CT Aβ+/NDAN-like (pBH = 0.00479) and Young versus Late AD (pBH = 0.0379). Significant 4-HNE contrasts were Young versus CT Aβ+/NDAN-like (pBH = 0.0314), Young versus Early AD (pBH = 0.00610) and Young versus Late AD (pBH = 0.0463). **D.** Median IHC-signature heatmap. Displayed raw medians were, for 4-HNE: 1.7, 2.3, 2.9, 3.5 and 2.8; GFAP: 0.3, 3.3, 8.3, 5.5 and 6.7; IBA1: 1.4, 7.0, 2.8, 9.8 and 4.7 across the same group order. **E.** Significant diagnosis-by-distance ligand-receptor interactions grouped by ligand-receptor pair: SPP1:CD44, 38 interactions; IL1B:IL1R1, 35; VEGFA:KDR, 4; TNF:TNFRSF1A, 1.

Underlying these cell-level associations, the spatial data indicate which signals carry between cells. Modeling ligand-receptor co-expression across distance and diagnosis, 78 diagnosis-by-distance interactions passed BH q < 0.05 (Supplemental Data Table S25). The osteopontin-CD44 gliosis pair SPP1:CD44 accounted for the largest and most consistent effects (38 interactions, all with positive diagnosis-by-distance coefficients, beta up to 0.013), identifying the astrocyte-to-microglia axis that couples the two glial populations. IL1B:IL1R1 interleukin-1 signaling appeared across as many cell pairs (35 interactions) but at much smaller magnitude (beta near 0.0001), with minor vascular VEGFA:KDR and TNF:TNFRSF1A contributions (**Fig. 7E**). These pairs name the communication routes, SPP1:CD44 in particular, through which the environment response is coordinated across the astrocytic, microglial and vascular-associated populations.

### 5. The same reactive-glia chemistry appears again across neurodegenerative diseases

The AD data place the inflammatory response in the tissue environment surrounding the plaque-rich region. If that response is a property of the environment, the same reactive chemistry should appear in other diseases that stress the same glial and myelin-rich tissue under a different trigger. We tested this in public single-nucleus datasets spanning multiple sclerosis (MS), amyotrophic lateral sclerosis (ALS) in motor cortex and in spinal cord [41], Parkinson disease (PD), Pick disease, progressive supranuclear palsy (PSP) and multiple system atrophy (MSA) [42–46]. In each dataset we scored the four AD-derived reactive-glia programs, iron, lipid-droplet LDAM, DAM-core and complement, in microglia and astrocytes against 1,000 expression-matched random gene sets, and separately tested whether the cell class that degenerates in that disease had lost redox and ferroptosis defense (Supplemental Data Table S26).

The lipid-droplet program was the most consistent signal. Microglial LDAM activity increased above the matched-null expectation in all seven disease-region tests, spanning MS, ALS motor cortex, ALS spinal cord, PD, Pick disease, PSP and MSA (**Fig. 8A, B**). The signal was strongest in MS white matter, where microglia carried all four reactive programs (LDAM permutation z = 9.9, p = 0.001; iron, DAM-core and complement all p ≤ 0.008). The other programs appeared in disease-specific combinations. ALS spinal cord carried microglial iron, LDAM and DAM-core alongside astrocytic iron, LDAM and complement. PD carried macrophage iron, LDAM and complement with astrocytic iron and DAM-core. Pick disease carried microglial LDAM, DAM-core and complement. MS gave the fullest and strongest response, matching the expectation that a white-matter myelin disease stresses the same tissue environment the AD data implicate.

The vulnerable cell in each disease showed the reciprocal change (**Fig. 8C**) Redox and ferroptosis defense fell in oligodendrocytes in MS white matter (delta = −0.220, matched-null z = −3.2, p = 0.003), in somatomotor neurons in ALS spinal cord (delta = −0.134, z = −4.4, p = 0.001), and in glutamatergic neurons in ALS motor cortex (delta = −0.041, z = −1.9, p = 0.042). The cell that degenerates in each disease is the cell losing its protection against the same lipid-peroxidation chemistry the reactive glia are engaging.

Across the seven datasets the trigger and the vulnerable cell change with the disease, and the reactive-glia chemistry stays the same. Reactive glia enter the lipid-droplet, iron and DAM-family states above matched nulls in every disease, and the vulnerable cell loses redox defense where the disease strikes hardest, in MS and ALS. This shows a convergence of signals of reactive environment, carbonyl stress and response, above matched nulls, not the AD program entirely. AD remains the one disease in this study where the same chemistry is also strongest away from the plaque, progresses with Braak stage, and its components move together as a self-reinforcing loop in the proteomic cohort. The cross-disease data place AD as the substrate-autonomous case of a reactive-glia chemistry that many catalysts can engage, which is what an environment-first, catalyst-agnostic reading of the disease predicts.

**Figure 8.**
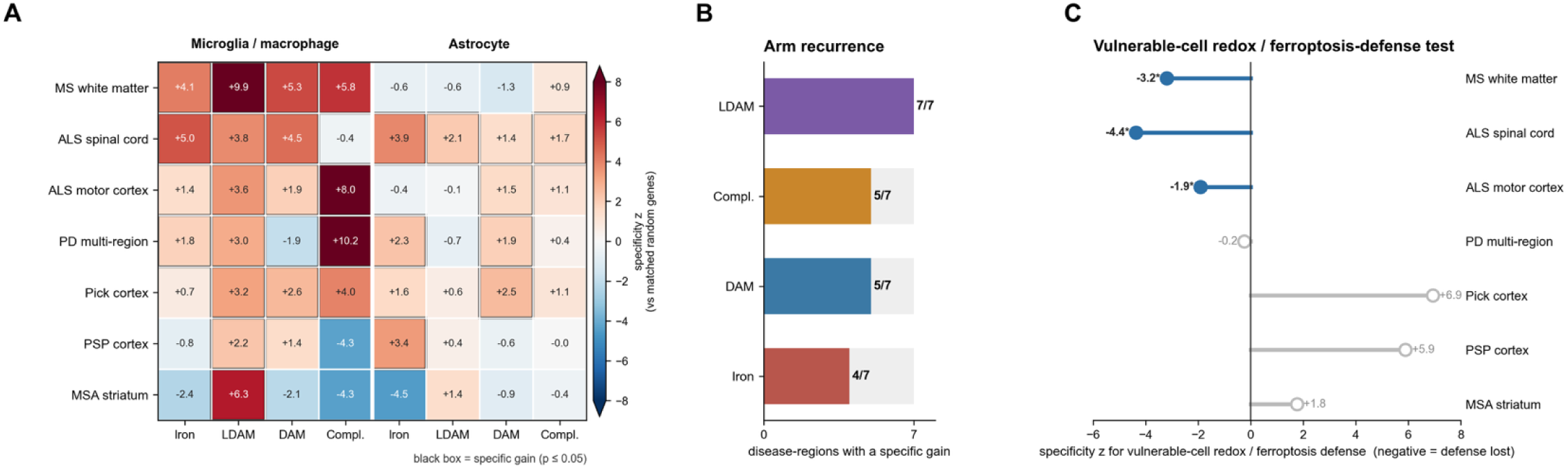
Single-cell cross-disease environment-state test. **A.** Reactive-glia specificity heatmap across seven disease-region single-nucleus datasets. Values show permutation z scores for disease-gained iron, LDAM, DAM-core and complement branches in microglia/macrophages and astrocytes against 1,000 expression-matched random gene sets. Black outlines mark specific gains, defined as empirical p <= 0.05 with positive permutation z and positive disease-control delta; 26 of 56 reactive cell-branch tests met this criterion. **B.** Branch frequency across disease-regions, counted once per disease-region when either reactive glial class showed a specific gain: LDAM 7/7, complement 5/7, DAM-core 5/7 and iron 4/7. **C.** Vulnerable-cell redox/ferroptosis-defense test, displaying the lowest-z vulnerable-cell test per disease-region. Negative z indicates defense loss and filled points mark significant losses: MS oligodendrocytes, z = -3.20, p = 0.002997; ALS spinal-cord somatomotor neurons, z = -4.37, p = 0.000999; ALS motor-cortex glutamatergic neurons, z = -1.92, p = 0.04196. Full single-cell results are provided in Supplemental Data Table S26.

## Discussion

Alzheimer’s disease is neuropathologically defined by its plaques and tangles, yet the inflammatory and chemical damage that correlate with cognitive symptoms are not the strongest where those lesions are densest. In our data, the strongest glial and oxidative reaction of the AD brain builds in the white matter and the myelin-rich, lipid-handling tissue around the plaques, while the amyloid itself accumulates in the neighboring grey matter, and the two maps separate. Plaque-associated gliosis is genuine and long described [6, 11, 12, 47–49], and yet most of the inflammatory chemistry we measure lies away from the plaque, inside a lipid-rich glial environment.

The amyloid cascade hypothesis places the generation and aggregation of Aβ at the start of a sequence that drives inflammation and synapse loss, leading to neurodegeneration [3, 4]. Our observations offer a different view of that sequence. The inflammatory response is anatomically dissociated from the plaques meant to drive it, and, in an independent single-nucleus cohort of several hundred donors, an oligodendrocyte lipid-droplet phenotype correlates with cognitive decline, even after accounting for amyloid and tangle burden [13]. Part of the tissue response therefore predicts cognitive decline beyond what the classical lesions explain. This gives spatial and molecular form to the cellular phase of AD: Aβ and pTau set off a tissue reaction that then propagates independently of the initiating lesions [50], and the modest, partial benefit of clearing amyloid in recent trials is consistent with this independent reaction [51, 52]. We propose that Aβ and pTau act as catalysts, tipping an already vulnerable tissue toward dementia, while the inflammatory response itself is carried by the environment they ignite.

We propose that the white-matter environment of the aging brain is metastable. Through life, it accumulates the substrate of its own vulnerability. It is the most PUFA-rich and iron-loaded compartment of the brain, it carries a constant metabolic demand for myelin turnover and lipid clearance, and oxidative and carbonyl modifications progressively accumulate with age [28–31, 53–55]. This is a primed but quiet condition, stable in the way a false vacuum is stable, holding until something lowers the barrier and lets it fall to a lower state where the catalyst tips the primed tissue into a self-reinforcing reaction. Three findings support the priming step on their own. In aged but cognitively healthy tissue, the white-matter lipid profile is already distinct, rich in myelin lipids and low in neutral storage fat, so the vulnerable chemistry is in place before plaques are dense. In aged Aβ-positive brains of people who remained cognitively healthy, microglial burden stays as low as in NDAN amyloid-negative controls [15, 16, 54]: the catalyst is present, but the tissue has not yet tipped into the reactive state. Across other diseases, the same environment enters a similar reaction under its own specific catalyst. Priming, catalyst and tipped state are each visible in separate cohorts. We hypothesize that primed brains progress to the tipped state, but this is a longitudinal claim that our cross-sectional data cannot settle on their own.

This tipped state has a defined chemistry: as Braak stage progresses, carbonyl and oxidative damage, AGE-RAGE and NF-κB signaling, iron handling and ferroptosis-related genes, lysosomal lipid disposal, complement and tissue remodeling all increase, with lipidomics showing cholesteryl esters and lysosomal lipids build-ups as myelin lipid is lost [18–20, 23, 28, 29, 35, 53–60]. We measured the direction of these changes and how they sharpen with disease stage, and we propose that they form a self-sustaining cycle. Iron-catalyzed peroxidation of the polyunsaturated lipids of myelin and glial membranes generates reactive carbonyls, such as 4-HNE. These carbonyls then drive AGE-RAGE and NF-κB signaling and impair lipid and lysosomal clearance, and the tissue left behind peroxidizes more readily than before [23, 24, 28–31, 61]. Each step of this loop is already established biochemistry, these data place the parts in the same compartment and show that they strengthen with disease stage, furthering the hypothesis of a self-reinforcing cycle. The reaction begins in white matter, the most peroxidizable and iron-loaded compartment of the brain [23, 28–31, 53–55]. The compartment that carries the strongest inflammatory gradient in these data is the one whose baseline chemistry is most ready to react.

Reactive glial phenotypes are well described, and several of them appear in the diseased brain: disease-associated microglia, lipid-droplet-accumulating microglia, disease-associated astrocytes, and the plaque-induced genes that surround Aβ and pTau deposits [6, 9, 35, 49, 56, 62–65]. These phenotypes are defined by their proximity to lesions, and by their gene expression changes. Our observations place those same reactive phenotypes in the environment far from the plaques and measure its lipid substrate directly by compartment-resolved lipidomics, so the chemistry is observed in the tissue itself.

The same chemistry also appears beyond AD. In single-nucleus data from multiple sclerosis, amyotrophic lateral sclerosis, Parkinson’s disease and cross-disorder dementia, the lipid-droplet reactive-glia state stands above expression-matched null in every disease-region test, and in each disease the vulnerable cell type loses its redox and ferroptosis defenses alongside it [42–46]. The catalyst and the vulnerable cell change from one disease to the next, while the chemistry of the reaction stays the same. While AD is the disease case we investigated most deeply through spatial transcriptomics, proteomic, lipidomic and immunohistochemical data, measurements from other datasets all converge on this self-sustaining inflammatory response following a catalyst.

Limitations. Spatial transcriptomics reports RNA from spot-level mixtures and does not resolve protein at the level of single cells. The lipidomics compares grey and white tissue within region by design, and its conclusions hold within region and compartment. The cross-disease analysis tests whether predefined response programs appear again in public datasets and is comparative support, not a causal experiment. We did not perturb the proposed redox-carbonyl cycle, so its closure within tissue and the direction of cause between substrate priming and reaction remain to be shown. We did not follow individual brains over time, so the passage from primed substrate to tipped state is read from cross-sectional contrasts.

These limitations set out the experiments that follow. If the substrate is primed before it tips, an aged, peroxidizable, iron-loaded white-matter chemistry should be detectable in cognitively healthy brains carrying little plaque, and measurable before symptoms begin. If the reaction runs with partial independence from amyloid, an intervention directed at redox and carbonyl chemistry should bend the trajectory of the tissue reaction without clearing amyloid, and the two approaches together should be testable against either alone. If the spot-level environmental signal reflects a cell-resolved state, single-nucleus and single-cell profiling of the same compartments should recover the oligodendrocyte and microglial states at cellular resolution. Each prediction can be tested, and each can fail, with methods already in hand or close to it.

## Conclusion

The inflammation in AD appears to be strongest in the lipid-rich glial and myelin environment that surrounds the plaques. It precedes them, and it runs as a redox-carbonyl reaction of failing lipid clearance, oxidative and carbonyl damage, glycation, iron mishandling and reactive gliosis. We propose that this environment is a metastable, primed state, and that Aβ and pTau are the catalysts that tip it toward dementia, with different catalysts tipping related environments in other neurodegenerative diseases. If the primed state feeds itself once tipped, it could outlast a falling plaque burden, and removing Aβ and pTau will not return the tissue to homeostatic conditions. Placing the primed environment at the centre of the disease, ahead of the catalyst that tips it, changes where the earliest pathology should be looked for and what a disease-modifying target has to reach.

## STAR Methods

### Resource availability

Lead contact. Requests for resources and further information should be directed to Aurélien M. Badina and Benjamin B. Tournier.

### Materials availability

This study did not generate new unique reagents.

### Data and code availability

Supplemental tables and figure source data cited in the manuscript are being finalized and are not included with this preprint. They, together with the analysis code, will be deposited in a citable public repository before journal submission. Raw human-tissue data are subject to donor-consent and institutional access restrictions.

### Experimental model and subject details

Human postmortem tissue was obtained through the Netherlands Brain Bank and Belle-Idée/local tissue resources under donor-consent and institutional tissue-use agreements.

#### Spatial cohort

16 FFPE hippocampal sections containing adjacent cortex (5 µm; CT 7, AD 9; ages 84-102; Braak I-VI; NBB). The full subregion-annotated QC-normalized spatial object contained 168,571 spots across hippocampal formation and adjacent cortex and was used for plaque burden, compartment-state, cortex/hippocampus slope, vascular/composition sensitivity and ligand-receptor analyses. A hippocampus-centered plaque-distance object used for gene-level plaque-distance discovery and module/carbonyl distance-curve analyses contained 84,389 spots; after full-metadata join, 80,606 matched hippocampal formation, 303 matched cortex and 3,480 did not carry a full subregion assignment. Aβ-positive spots were defined from manual anti-Aβ42 ROI annotation and propagated as plaque-covered spots, boolean plaque status and categorical Ab+/Ab-status. Fig. 1A used the full QC-retained spot denominator grouped per section.

#### Proteomic cohort

31 fresh-frozen hippocampal samples (CT 6, EAD 12, LAD 13; Braak II–VI).

#### Lipidomics cohort

94 submitted grey/white tissue samples from cortex and hippocampus, with 90 analyzed positive-mode lipidomics samples from 49 subjects; NBB NG/NW samples were analyzed as cortex, Belle-Idée G/W samples as hippocampus, and grey/white comparisons were performed within region. These cohorts are analyzed as separate human-tissue resources unless a specific identifier-matched analysis is stated.

#### FFPE-IHC cohort

In the paired spatial cohort, complete GML/WML measurements were available for 5 CT and 10 AD subjects for AT8, 4G8, GFAP and IBA1. For IBA1, the samples contained one CT and one AD subject represented only in the mixed compartment, and we excluded them from paired GML/WML contrasts. The independent source cohort comprised 54 subjects spanning young, CT Aβ−, CT Aβ+/NDAN-like, EAD and LAD groups. Before outlier filtering, the imported marker tables contained 51 IBA1, 70 GFAP and 47 4-HNE measurements. Tukey 1.5 × IQR filtering within marker × diagnostic group excluded 2 IBA1 and 6 GFAP measurements and no 4-HNE measurements, leaving 49, 64 and 47 measurements, respectively.

### Methods

#### Spatial transcriptomics

10x Genomics Visium CytAssist with immunofluorescence pre-processing; Space Ranger v3.0.1, GRCh38-2020-A; Seurat v5; SCTransform + Harmony for the hippocampus-centered plaque-distance object, and log-normalized full-object analyses for subregion-resolved cortex/hippocampus tests. Manual ROI annotations defined hippocampal formation subregions, adjacent cortex, GM/GML and WM/WML compartments and Aβ-positive spots. Plaque distance was computed as the nearest-neighbor Euclidean distance from each spot to the closest Aβ-positive spot, scaled by Space Ranger pixel size. Gene-level plaque-distance models used 100 µm slide-level shell pseudobulks over 0 <= distance < 1000 µm, raw Spatial counts, voom-quality weights and duplicate correlation by slide. Fig. 2C module curves used exact spot-level distances in AD sections over 0 < distance <= 900 µm and plotted shell midpoints from 50 to 850 µm; Aβ-positive spots at distance 0 and all spots >900 µm were excluded.

#### RCTD spot deconvolution

Green et al. 2024 ROSMAP dorsolateral prefrontal cortex (DLPFC) sketch atlas reference (45,577 nuclei, 17 broad classes, 95 fine states); per-slide doublet mode for broad classes, full mode for fine states. Collapsed-7 broad-class adjustment for composition rescue. Vascular/BBB sensitivity analyses used shell-level weighted linear models with SlideID fixed effects: the far-from-plaque inflammatory-gradient aggregate was adjusted for RCTD endothelial, vascular/mural and erythrocyte weights, and canonical SCT marker scores were constructed as mean z-scored expression across endothelial/BBB, mural/perivascular and blood-leakage genes before WML plaque-distance models.

#### Bulk DIA proteomics

Tris-NaCl-Triton lysis, sonication, 20,000 g 20 min 4 °C; trypsin digestion; nanoLC-MS/MS Vanquish NEO + Orbitrap Fusion Lumos DIA mode; directDIA+ (Spectronaut) at 1% FDR with ≥ 2 unique peptides per protein. 5,422 proteins quantified across 31 subjects.

#### Cross-disease single-cell environment-state scoring

public single-nucleus datasets were analyzed at cell level for MS white matter, ALS motor cortex, ALS spinal cord, PD multi-region brain, Rexach PSP / Pick cortical tauopathy and MSA striatum. Counts were streamed from backed AnnData objects, normalized per cell as CP10K and log1p transformed. For each dataset and cell type, response-branch scores were computed as mean expression of the gene set minus the cell’s mean expression across the scored/background gene pool. Reactive glia were tested for disease-control increases in four disease-gained response branches: iron (FTL, FTH1), lipid-droplet / LDAM (PLIN2, PLIN3, APOE, GPNMB, LPL, FABP5, SOAT1, ACSL1, DGAT2), DAM-core (TREM2, TYROBP, CST7, CD68, ITGAX, LGALS3, SPP1, CD44) and complement (C1QA, C1QB, C1QC, C3). Specificity was measured against 1,000 random expression-matched gene sets of equal size, sampled from 25 expression bins; empirical p-values used the upper tail for reactive-glia gain. Vulnerable-cell redox/ferroptosis defense was tested with GPX4, ACSL4, SLC7A11, NCOA4, SOD1, PRDX1, FTH1, FTL, HSPA1A and DNAJB1, using one-sided Mann-Whitney disease-control decrease and the lower tail of the same expression-matched null. Housekeeping and ribosomal genes were not used as controls because ribosome biogenesis can itself be part of cellular activation. PD vulnerable-cell analysis used GABAergic neurons as a broad-label proxy because the local Roussos H5AD did not expose a dopaminergic-neuron label.

#### Gradient-module bulk proteomic composite and module-level Braak models

49 source-annotated inflammation modules were scored in spatial data and hippocampal proteomics. The 116-gene inflammatory-gradient core was defined as the union of 13 modules with WML-far/GML-near topology at module-score level. For proteomics, per-subject module abundance was the mean log2 PG-quantity across detected proteins for modules with at least five detected proteins. The proteomic inflammatory-gradient composite was the per-subject z-score averaged across the nine bulk-proteomics-testable spatial-gradient modules.

#### Gene-level audit and Supplemental Data Table S16

all available pathway gene integrated summaries were scanned for genes with both spatial plaque-distance evidence and proteomic Braak evidence. Benjamini-Hochberg q-values were recomputed across the scan for spatial all-subject plaque-distance p-values and proteomic Braak p-values. Candidate genes were defined by spatial scan q < 0.05 and proteomic Braak scan q < 0.20, then deduplicated by gene using the combined spatial/proteomic rank. Direction classes were assigned from the AD spatial distance slope and the proteomic Braak slope. Raw-count traceability was assessed from the raw Spatial assay as log1p(CP10K) after aligning spots to the distance metadata. The 29 pathway overlays were locked before manuscript drafting from WikiPathways, KEGG and MSigDB Hallmark gene sets, then filtered to AD-relevant carbonyl/redox, AGE-RAGE/NF-κB, lipid-peroxidation/ferroptosis, proteostasis, vascular/BBB, synaptic and glial-response biology. Pathway-overlay summaries used the pathway_gene_map_and_status tables; in these tables, spatial_delta_all_near_far > 0 means higher near plaques and spatial_delta_all_near_far < 0 means higher far from plaques.

#### Carbonyl/redox compartment-signature scoring

the 31-gene carbonyl-stress core was assembled from glyoxalase/MGO detoxification, fructosamine repair, aldo-keto/aldehyde carbonyl clearance, NRF2-glutathione antioxidant response and peroxide/lipid-peroxidation detoxification genes. Six broader signatures were assembled for AGE-RAGE ligand biology, MGO/glyoxalase detoxification, glutathione/NRF2 redox response, lipid-peroxidation/ferroptosis, aldehyde-carbonyl clearance and protein-damage/proteostasis. For each signature, detected genes were averaged per slide x compartment; paired WML-WM, GML-GM and WML-GML contrasts used two-sided paired Wilcoxon signed-rank tests with BH correction.

#### Positive-mode RPLC-HRMS lipidomics

lipidomics was analyzed as compositional data. Raw lipid species intensities were joined to the analyzed sample metadata, mapped to cortex or hippocampus by provenance and summarized within grey and white compartments. Class and ratio scores were computed using CLR/log-ratio transformations instead of total abundance. CT, EAD and LAD contrasts were tested within region × compartment using two-sided rank-based tests and Braak Spearman correlations, followed by Benjamini-Hochberg correction within the lipid-class family. Species-level cortex-white LAD-versus-CT tests used the same within-compartment framework. Lipidomics-proteomics overlap analyses used matched subject identifiers and the reduced eight-anchor proteomic loop PC1.

#### Green atlas module scoring

the 49 inflammation modules were scored on the 45,577-nucleus sketch reference derived from the Green et al. 2024 ROSMAP dorsolateral prefrontal cortex (DLPFC) atlas [13]. Per-cell log-normalized module scores were aggregated by donor and broad cell type. Cognitive-decline models were fit per module × cell-type family as cog_slope ∼ module + age + sex + sqrt-amyloid + sqrt-tangles. The model set included microglia, astrocytes, oligodendrocytes, endothelial cells, CUX2+ excitatory neurons and inhibitory neurons. Benjamini-Hochberg correction was applied within each cell-type family.

#### Immunohistochemistry

spatial-cohort 5 µm FFPE sections were stained for AT8 (anti-phospho-tau, Thermo MN1020), 4G8 (anti-Aβ, BioLegend SIG-39220), GFAP (rabbit polyclonal, Dako Z0334) and IBA1 (rabbit polyclonal, Wako 019-19741); DAB visualisation; QuPath quantification per region × compartment by a blinded operator. The spatial-cohort subregional supplement reports the raw subject × region × compartment %-positive-area values. The independent 54-subject FFPE cohort (12 µm Belle Idée sections) was analyzed with outlier-aware group summaries and Kruskal-Wallis / Dunn or Wilcoxon rank-sum post-hoc tests for IBA1, GFAP and 4-HNE.

#### Cross-disease supplemental reporting

disease-region single-nucleus reactive and vulnerable-cell tests, permutation z-scores, empirical p-values and cell counts are provided in Supplemental Data Table S26. Per-disease results, negative-control corrections, specificity screens and dataset inventories are provided in Supplemental Data Tables S27-S36.

### Quantification and statistical analysis

All multiple-testing corrections used Benjamini-Hochberg q-values within each analysis and modality family. Primary manuscript significance is reported at q <= 0.05, with exact p and q values retained in supplemental tables; q < 0.20 is used only for explicitly exploratory screens. Human-tissue models included sex, age and PMD as covariates when available. Mixed-effects models used lme4/nlme REML with subject-level or slide-level random intercepts where applicable. Paired compartment contrasts used two-sided paired Wilcoxon signed-rank tests. Group contrasts used two-sided rank-based tests unless stated otherwise. Spatial ligand-receptor proxy models fit mean_score ∼ dist100 * diagnosis + region + compartment + plaque-burden z-score, with slide-level random intercepts where applicable.

## CRediT author statement

**A.M.B.**: Conceptualization, Methodology, Software, Validation, Formal Analysis, Investigation, Data Curation, Writing - Original Draft, Writing - Review & Editing, Visualization, Supervision, Project Administration. **E.R.**: Formal analysis, Writing - Review & Editing. **I.M.**: Methodology, Formal analysis, Writing - Review & Editing. **Q.A.**: Formal analysis, Investigation, Writing - Review & Editing. **L.A.**: Writing - Review & Editing. **K.C.**: Writing - Review & Editing. **S.T.**: Conceptualization, Writing - Review & Editing. **S.R.**: Methodology, Writing - Review & Editing. **P.M.**: Conceptualization, Resources, Writing - Review & Editing. **B.B.T.**: Conceptualization, Validation, Formal analysis, Investigation, Resources, Supervision, Project Administration, Funding Acquisition, Writing - Original Draft, Writing - Review & Editing.

## Acknowledgments

We thank Pia Lovero for the technical assistance with the immunohistochemistry. We thank the Netherlands Brain Bank for tissue donation and clinical metadata. We thank Mylène Docquier, Didier Chollet and the iGE3 Genomics Platform (University of Geneva) for their sequencing and support, and Alexandre Hainard and Domitille Schvartz and the UNIGE Proteomics Core Facility for their mass-spectrometry support. We also thank Gilad Sahar Green and Vilas Menon for the public release of the ROSMAP DLPFC single-nucleus reference atlas.

## References

1. Montine, T.J., et al., National Institute on Aging-Alzheimer’s Association guidelines for the neuropathologic assessment of Alzheimer’s disease: a practical approach. Acta Neuropathol, 2012. 123(1): p. 1–11.

2. Braak, H. and E. Braak, Neuropathological stageing of Alzheimer-related changes. Acta Neuropathol, 1991. 82(4): p. 239–59.

3. Hardy, J.A. and G.A. Higgins, Alzheimer’s disease: the amyloid cascade hypothesis. Science, 1992. 256(5054): p. 184–5.

4. Hardy, J. and D.J. Selkoe, The amyloid hypothesis of Alzheimer’s disease: progress and problems on the road to therapeutics. Science, 2002. 297(5580): p. 353–6.

5. Heneka, M.T., et al., Neuroinflammation in Alzheimer’s disease. Lancet Neurol, 2015. 14(4): p. 388– 405.

6. Keren-Shaul, H., et al., A Unique Microglia Type Associated with Restricting Development of Alzheimer’s Disease. Cell, 2017. 169(7): p. 1276–1290 e17.

7. Krasemann, S., et al., The TREM2-APOE Pathway Drives the Transcriptional Phenotype of Dysfunctional Microglia in Neurodegenerative Diseases. Immunity, 2017. 47(3): p. 566–581 e9.

8. Mathys, H., et al., Single-cell transcriptomic analysis of Alzheimer’s disease. Nature, 2019. 570(7761): p. 332–337.

9. Habib, N., et al., Disease-associated astrocytes in Alzheimer’s disease and aging. Nat Neurosci, 2020. 23(6): p. 701–706.

10. Zhou, Y., et al., Human and mouse single-nucleus transcriptomics reveal TREM2-dependent and TREM2-independent cellular responses in Alzheimer’s disease. Nat Med, 2020. 26(1): p. 131–142.

11. Gerrits, E., et al., Distinct amyloid-beta and tau-associated microglia profiles in Alzheimer’s disease. Acta Neuropathol, 2021. 141(5): p. 681–696.

12. Huang, Y., et al., Regulation of cell distancing in peri-plaque glial nets by Plexin-B1 affects glial activation and amyloid compaction in Alzheimer’s disease. Nat Neurosci, 2024. 27(8): p. 1489–1504.

13. Green, G.S., et al., Cellular communities reveal trajectories of brain ageing and Alzheimer’s disease. Nature, 2024. 633(8030): p. 634–645.

14. Mathys, H., et al., Single-cell atlas reveals correlates of high cognitive function, dementia, and resilience to Alzheimer’s disease pathology. Cell, 2023. 186(20): p. 4365–4385 e27.

15. Iliff, J.J., et al., A paravascular pathway facilitates CSF flow through the brain parenchyma and the clearance of interstitial solutes, including amyloid beta. Sci Transl Med, 2012. 4(147): p. 147ra111.

16. Simon, M.J. and J.J. Iliff, Regulation of cerebrospinal fluid (CSF) flow in neurodegenerative, neurovascular and neuroinflammatory disease. Biochim Biophys Acta, 2016. 1862(3): p. 442–51.

17. O’Brien, J.S. and E.L. Sampson, Fatty acid and fatty aldehyde composition of the major brain lipids in normal human gray matter, white matter, and myelin. J Lipid Res, 1965. 6(4): p. 545–51.

18. Sayre, L.M., et al., 4-Hydroxynonenal-derived advanced lipid peroxidation end products are increased in Alzheimer’s disease. J Neurochem, 1997. 68(5): p. 2092–7.

19. Williams, T.I., et al., Increased levels of 4-hydroxynonenal and acrolein, neurotoxic markers of lipid peroxidation, in the brain in Mild Cognitive Impairment and early Alzheimer’s disease. Neurobiol Aging, 2006. 27(8): p. 1094–9.

20. Deane, R., et al., RAGE mediates amyloid-beta peptide transport across the blood-brain barrier and accumulation in brain. Nat Med, 2003. 9(7): p. 907–13.

21. Yan, S.D., et al., RAGE and amyloid-beta peptide neurotoxicity in Alzheimer’s disease. Nature, 1996. 382(6593): p. 685–91.

22. Markesbery, W.R. and M.A. Lovell, Four-hydroxynonenal, a product of lipid peroxidation, is increased in the brain in Alzheimer’s disease. Neurobiol Aging, 1998. 19(1): p. 33–6.

23. Ashraf, A., et al., Iron dyshomeostasis, lipid peroxidation and perturbed expression of cystine/glutamate antiporter in Alzheimer’s disease: Evidence of ferroptosis. Redox Biol, 2020. 32: p. 101494.

24. Smith, M.A., et al., Iron accumulation in Alzheimer disease is a source of redox-generated free radicals. Proc Natl Acad Sci U S A, 1997. 94(18): p. 9866–8.

25. Huang, X., et al., Cu(II) potentiation of alzheimer abeta neurotoxicity. Correlation with cell-free hydrogen peroxide production and metal reduction. J Biol Chem, 1999. 274(52): p. 37111–6.

26. Curtain, C.C., et al., Alzheimer’s disease amyloid-beta binds copper and zinc to generate an allosterically ordered membrane-penetrating structure containing superoxide dismutase-like subunits. J Biol Chem, 2001. 276(23): p. 20466–73.

27. Pratico, D., et al., Increased F2-isoprostanes in Alzheimer’s disease: evidence for enhanced lipid peroxidation in vivo. FASEB J, 1998. 12(15): p. 1777–83.

28. Dixon, S.J., et al., Ferroptosis: an iron-dependent form of nonapoptotic cell death. Cell, 2012. 149(5): p. 1060–72.

29. Hambright, W.S., et al., Ablation of ferroptosis regulator glutathione peroxidase 4 in forebrain neurons promotes cognitive impairment and neurodegeneration. Redox Biol, 2017. 12: p. 8–17.

30. Ayton, S., et al., Ferritin levels in the cerebrospinal fluid predict Alzheimer’s disease outcomes and are regulated by APOE. Nat Commun, 2015. 6: p. 6760.

31. Bao, W.D., et al., Loss of ferroportin induces memory impairment by promoting ferroptosis in Alzheimer’s disease. Cell Death Differ, 2021. 28(5): p. 1548–1562.

32. Sanchez-Mejia, R.O., et al., Phospholipase A2 reduction ameliorates cognitive deficits in a mouse model of Alzheimer’s disease. Nat Neurosci, 2008. 11(11): p. 1311–8.

33. Ioannou, M.S., et al., Neuron-Astrocyte Metabolic Coupling Protects against Activity-Induced Fatty Acid Toxicity. Cell, 2019. 177(6): p. 1522–1535 e14.

34. Shi, Y., et al., ApoE4 markedly exacerbates tau-mediated neurodegeneration in a mouse model of tauopathy. Nature, 2017. 549(7673): p. 523–527.

35. Marschallinger, J., et al., Lipid-droplet-accumulating microglia represent a dysfunctional and proinflammatory state in the aging brain. Nat Neurosci, 2020. 23(2): p. 194–208.

36. Haney, M.S., et al., APOE4/4 is linked to damaging lipid droplets in Alzheimer’s disease microglia. Nature, 2024. 628(8006): p. 154–161.

37. Blanchard, J.W., et al., APOE4 impairs myelination via cholesterol dysregulation in oligodendrocytes. Nature, 2022. 611(7937): p. 769–779.

38. Latimer, C.S., et al., Resistance and resilience to Alzheimer’s disease pathology are associated with reduced cortical pTau and absence of limbic-predominant age-related TDP-43 encephalopathy in a community-based cohort. Acta Neuropathol Commun, 2019. 7(1): p. 91.

39. Zolochevska, O. and G. Taglialatela, Non-Demented Individuals with Alzheimer’s Disease Neuropathology: Resistance to Cognitive Decline May Reveal New Treatment Strategies. Curr Pharm Des, 2016. 22(26): p. 4063–8.

40. Perez-Nievas, B.G., et al., Dissecting phenotypic traits linked to human resilience to Alzheimer’s pathology. Brain, 2013. 136(Pt 8): p. 2510–26.

41. Takeuchi, E., et al., Single-nucleus multiome shows motor neuron glutamate overactivation in amyotrophic lateral sclerosis. Brain, 2026. 149(7): p. 2480–2494.

42. Hametner, S., et al., Iron and neurodegeneration in the multiple sclerosis brain. Ann Neurol, 2013. 74(6): p. 848–61.

43. Absinta, M., et al., A lymphocyte-microglia-astrocyte axis in chronic active multiple sclerosis. Nature, 2021. 597(7878): p. 709–714.

44. Jakel, S., et al., Altered human oligodendrocyte heterogeneity in multiple sclerosis. Nature, 2019. 566(7745): p. 543–547.

45. Kamath, T., et al., Single-cell genomic profiling of human dopamine neurons identifies a population that selectively degenerates in Parkinson’s disease. Nat Neurosci, 2022. 25(5): p. 588–595.

46. Rexach, J.E., et al., Cross-disorder and disease-specific pathways in dementia revealed by single-cell genomics. Cell, 2024. 187(20): p. 5753–5774 e28.

47. Miyoshi, E., et al., Spatial and single-nucleus transcriptomic analysis of genetic and sporadic forms of Alzheimer’s disease. Nat Genet, 2024. 56(12): p. 2704–2717.

48. Avey, D.R., et al., Uncovering plaque-glia niches in human Alzheimer’s disease brains using spatial transcriptomics. Mol Neurodegener Adv, 2025. 1(1): p. 2.

49. Chen, W.T., et al., Spatial Transcriptomics and In Situ Sequencing to Study Alzheimer’s Disease. Cell, 2020. 182(4): p. 976–991 e19.

50. De Strooper, B. and E. Karran, The Cellular Phase of Alzheimer’s Disease. Cell, 2016. 164(4): p. 603– 15.

51. van Dyck, C.H., et al., Lecanemab in Early Alzheimer’s Disease. N Engl J Med, 2023. 388(1): p. 9–21.

52. Sims, J.R., et al., Donanemab in Early Symptomatic Alzheimer Disease: The TRAILBLAZER-ALZ 2 Randomized Clinical Trial. JAMA, 2023. 330(6): p. 512–527.

53. Han, X., et al., Substantial sulfatide deficiency and ceramide elevation in very early Alzheimer’s disease: potential role in disease pathogenesis. J Neurochem, 2002. 82(4): p. 809–18.

54. Han, X., Potential mechanisms contributing to sulfatide depletion at the earliest clinically recognizable stage of Alzheimer’s disease: a tale of shotgun lipidomics. J Neurochem, 2007. 103 Suppl 1(Suppl 1): p. 171–9.

55. Depp, C., et al., Myelin dysfunction drives amyloid-beta deposition in models of Alzheimer’s disease. Nature, 2023. 618(7964): p. 349–357.

56. Xu, Z., et al., Microglia-specific regulation of lipid metabolism in Alzheimer’s disease revealed by microglial depletion in 5xFAD Mice. Nat Commun, 2025. 16(1): p. 9156.

57. Heneka, M.T., et al., NLRP3 is activated in Alzheimer’s disease and contributes to pathology in APP/PS1 mice. Nature, 2013. 493(7434): p. 674–8.

58. Ising, C., et al., NLRP3 inflammasome activation drives tau pathology. Nature, 2019. 575(7784): p. 669–673.

59. Hong, S., et al., Complement and microglia mediate early synapse loss in Alzheimer mouse models. Science, 2016. 352(6286): p. 712–716.

60. van der Kant, R., et al., Cholesterol Metabolism Is a Druggable Axis that Independently Regulates Tau and Amyloid-beta in iPSC-Derived Alzheimer’s Disease Neurons. Cell Stem Cell, 2019. 24(3): p. 363– 375 e9.

61. Nixon, R.A., et al., Extensive involvement of autophagy in Alzheimer disease: an immuno-electron microscopy study. J Neuropathol Exp Neurol, 2005. 64(2): p. 113–22.

62. Nugent, A.A., et al., TREM2 Regulates Microglial Cholesterol Metabolism upon Chronic Phagocytic Challenge. Neuron, 2020. 105(5): p. 837–854 e9.

63. Gouna, G., et al., TREM2-dependent lipid droplet biogenesis in phagocytes is required for remyelination. J Exp Med, 2021. 218(10).

64. Liddelow, S.A., et al., Neurotoxic reactive astrocytes are induced by activated microglia. Nature, 2017. 541(7638): p. 481–487.

65. Jo, S., et al., GABA from reactive astrocytes impairs memory in mouse models of Alzheimer’s disease. Nat Med, 2014. 20(8): p. 886–96.

